# Hydrophobic-cationic peptides enhance RNA polymerase ribozyme activity by accretion

**DOI:** 10.1101/2021.02.22.432394

**Authors:** Peiying Li, Philipp Holliger, Shunsuke Tagami

**Author notes:** Corresponding Authors: P.H, S.T.

## Abstract

Accretion and the resulting increase in local concentration to enhance target stability and function is a widespread mechanism in biology (for example in the liquid-liquid demixing phases and coacervates). It is widely believed that such macromolecular aggregates (formed through ionic and hydrophobic interactions) may have played a role in the origin of life. Here, we report on the behaviour of a hydrophobic-cationic RNA binding peptide selected by phage display (P43: AKKVWIIMGGS) that forms insoluble aggregates, accrete RNA on their surfaces in a size-dependent manner, and thus enhance the activities of various ribozymes. At low Mg^2+^ concentrations ([Mg^2+^]: 25 mM MgCl_2_), the activity of a small ribozyme (hammerhead ribozyme) was enhanced by P43, while larger ribozymes (RNA polymerase ribozyme (RPR), RNase P, F1* ligase) were inhibited. In contrast, at high [Mg^2+^] (≥200 mM), the RPR activity was enhanced. Another hydrophobic-cationic peptide with a simpler sequence (K_2_V_6_: KKVVVVVV) also exhibited similar regulatory effects on the RPR activity. Furthermore, inactive RPR captured on P43 aggregates at low [Mg^2+^] could be reactivated in a high [Mg^2+^] buffer. Therefore, in marked contrast to previously studied purely cationic peptides (like K_10_) that enhance RPR only at low ionic strength, hydrophobic-cationic peptides can reversibly concentrate RNA and enhance the RPR activity even at high ionic strength conditions such as in eutectic ice phases. Such peptides could have aided the emergence of longer and functional RNAs in a fluctuating environment (e.g., dry-wet / freeze-thaw cycles) on the prebiotic earth.

## INTRODUCTION

Accretion and self-organization of biopolymers (e.g., peptides and RNA) into defined locations were likely important steps in the emergence of life. Indeed, compartmentalization or spatial confinement are seen as particularly important for the emergence of genetic self-replication and Darwinian evolution, as otherwise genetic kin selection cannot occur and a replication system becomes vulnerable to be overrun by replication parasites.^1,2^ In all modern organisms, cell membranes made of phospholipids and, to some extent, polypeptide scaffolds and demixing phases maintain the spatial organization and integration of the complex macromolecular systems. However, how in what form such organizing principles emerged on the prebiotic earth and what molecules/structures achieved accumulation of functional biopolymers has remained elusive. As the non-biological synthesis of phospholipids is relatively complicated, simpler fatty-acid membranes have also been suggested to have formed ancient cell-like structures.^3,4^ However, they are unstable in conditions containing divalent cations (e.g., Mg^2+^), which are, in turn, thought to be necessary to support structures and functions of RNA.^5–7^ Considering RNA is widely believed to have been a pivotal molecule for the primitive replication system,^8,9^ such a favorable environment also needs to be compatible with ribozyme, in particular, RNA polymerase ribozyme (RPR) to drive RNA replication.^10,11^ Although citrate can reduce the destabilization of fatty acid membranes by divalent cations,^12^ it was found to strongly inhibit the RPR activity.^13^

Peptides are another type of biopolymer that could have existed on the prebiotic earth.^14–17^ We previously showed that simple cationic peptides (e.g., oligolysine) could reduce the strong Mg^2+^ dependency of RPR likely by enhancing interactions between negatively-charged RNA molecules promoting assembly of the RPR holoenzyme.^13^ Oligolysine also stimulates the activity of the RNase P ribozyme, an ancient ribozyme conserved in all three domains of life,^13,18^ indicating cationic peptides might have functioned as general co-factors for many ribozymes. Various plausible pathways for the prebiotic synthesis of such charged peptides have been suggested.^14,15^ For example, thermodynamic calculation predicted the prebiotic synthesis of positively/negatively-charged peptides on mineral surfaces.^19,20^ Depsipeptides containing cationic proteinaceous amino acid residues (Lys, Arg) can be synthesized in prebiotically plausible dry-down reactions and mutually stabilize RNA.^21,22^ All of these empirical observations together with theoretical considerations (as well as analogies with extant biology) support the idea that simple peptides could have aided the assembly and function of RNA-based biosystems.

Furthermore, peptides can spontaneously form aggregates and liquid-liquid phase-separations (LLPSs) in modern cells and test tubes, providing another candidate pathway for the formation of prebiotic biomolecular assemblies.^16,17, 23–25^ Indeed, the hammerhead ribozyme (∼30 nt) can perform its self-cleavage reaction in LLPSs containing positively-charged peptides (e.g., oligolysine),^26,27^ and its activity is enhanced by addition of a negatively charged peptide (D_10_) presumably by loosening too tight RNA-polycation interactions.^28^ However, the formation of such peptide-RNA LLPSs requires unrealistically high concentrations of both peptides and RNAs on the prebiotic earth. Alternatively, peptide assemblies could have formed more easily, as peptides can be synthesized in various prebiotic conditions^14,15^ and they spontaneously assemble forming insoluble aggregates (e.g., β-amyloid), which may have formed a scaffold for other molecules to organize on and even could have self-propagated.^17^ For example, simple peptides with alternating hydrophobic and hydrophilic residues (e.g., Val-Lys repeats) can form amyloid structures,^17,29^ and can also be synthesized in prebiotic chemistry by using peptides with complementary charges as templates.^30^ Furthermore, such peptides and nucleic acids can mutually promote amyloid formation and hybridization.^31^ However, the effect of such peptide aggregates on ribozyme functions has not been studied in detail yet.

Here, we report an RNA-binding peptide de novo selected by phage display (P43: AKKVWIIMGGS) for binding to an RPR construct, Zc.^32^ P43 forms insoluble aggregates and shows both stimulative and inhibitive effects on various ribozymes depending on ribozyme size, ionic strength and Mg^2+^ concentration ([Mg^2+^]). We characterize these effects and find that P43 - unlike oligolysine peptides function only at low [Mg^2+^] and therefore would lose their effect in highly concentrated solutions such as eutectic water ice phases^13^-can concentrate RNAs and stimulate RPR activity at high [Mg^2+^]. Such regulations of the RPR activity could also be performed by a simpler hydrophobic-cationic peptide (K_2_V_6_: KKVVVVVV). We argue that such hydrophobic-cationic peptide aggregates could have captured diluted nucleic acids at the wet phases and activated ribozymes at nearly-dry or frozen phases of the hypothetical tidal wet-dry or diurnal freeze-thaw cycles, to accumulate and select functional RNA molecules.

## RESULTS AND DISCUSSION

### Selection of peptides to regulate the RPR activity

Aiming to obtain new peptides that bind and regulate an RNA polymerase ribozyme (RPR Z^LT^, Figure 1A), we first performed peptide selection by applying the previously-reported phage display protocol.^33^ We prepared a phagemid library with seven randomized amino acid residues (AXXXXXXXGGS) and also synthesized a biotinylated RPR construct stabilized in the active conformation as the target molecule (RPR Zc, Supplementary Figure 1A).^13,32^ Then, we performed 3–4 rounds of selection of the peptide library against the ribozyme immobilized on magnetic beads and sequenced isolated phagemids to identify the selected peptide sequences (Supplementary Figure 1B). Roughly 70% of the selected sequences contained a common motif (KCCF/Y), and no other sequence motif was identified. Then, we tested the effects of twelve selected peptides (three with the KCCF/Y motif and nine without it) on the activity of RPR in a condition containing 25 mM MgCl_2_ (Figure 1B). While most of the peptide did not show any significant effect, a hydrophobic-cationic peptide (P43: AKKVWIIMGGS) strongly inhibited the activity of RPR. We also found P43 did not work at low concentration (50–200 µM), while relatively large amounts of P43 (≥ 400 µM) almost completely inhibited the activity of RPR Z^LT^ (Figure 1C, D).

**Figure 1.**
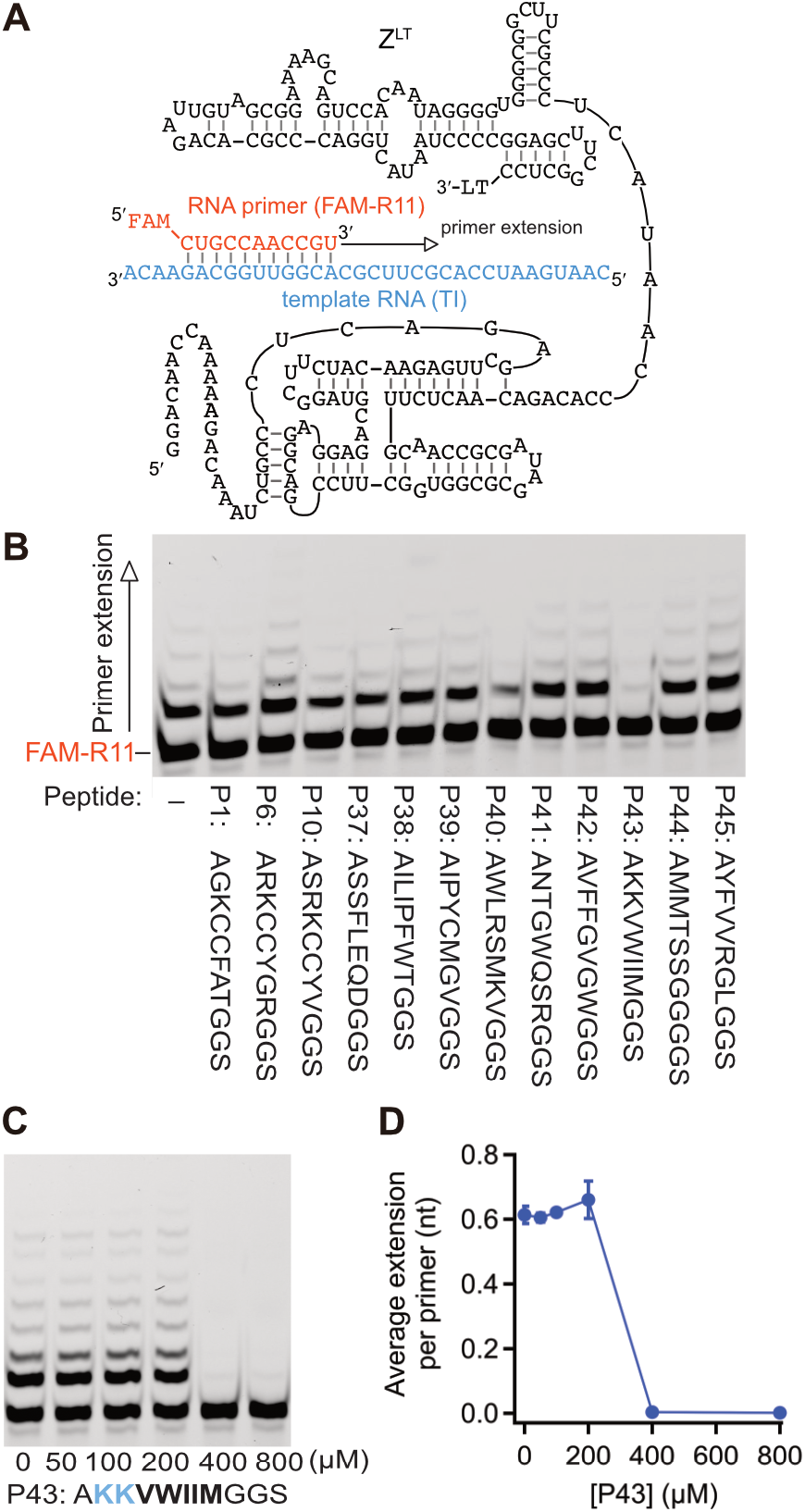
Selection of ribozyme-binding peptides. (A) Schematic depiction of an RNA polymerase ribozyme (RPR Z^LT^). 3′-LT is a 3′-extension sequence (GCGGCCGCAAAAAAAAAAGGCUUACC) used in the previous ribozyme selection for RPR Z.^11^ (B) Primer extension by RPR Z^LT^ was performed in 50 mM Tris•HCl ((B) pH 7.6, (C, D) pH 8.3) buffer containing 25 mM MgCl_2_, 8% PEG6000, 0.4%DMSO and 0.5 mM of each NTP. The reactions were incubated at 17°C for 7 days. (A) Concentration of the peptides was 0.4 mg/mL except for P42, which was not soluble and used as a 0.04 mg/mL suspension. (D) Error bars represent S.D. (N = 3).

### P43 forms insoluble aggregates and captures RNA

As P43 functioned only when its concentration was high (Figure 1C, D), we speculated that P43 would have worked by forming aggregates and carefully examined its stock solution. When the peptide was dissolved in DMSO and diluted with water, the solution (20 mg/mL P43, 20% DMSO) was apparently clear. However, after keeping it in a freezer overnight or longer, soft opaque precipitations were formed (Supplementary Figure 2). Thus, we supposed the P43 peptide functions as RNA-binding aggregates. The resuspended P43 aggregates showed the typical CD spectra for random coil peptides with a negative peak at 198 nm (Supplementary Figure 3A). Electron microscopic analysis revealed the P43 sample contains aggregates of various forms (Supplementary Figure 4A–D). However, when we shot X-ray on the P43 aggregates, we could observe X-ray reflections at 4.7 Å and 11.5 Å (Supplementary Figure 5), which indicates the aggregates at least partially adopt the cross-β conformation. Interestingly, in the presence of NTPs, another negative peak at 215 nm appeared in the CD spectra of P43 (Supplementary Figure 3A). As only ∼10% (0.05 mM) of the mixed NTPs were estimated to co-precipitated with P43 (Supplementary Figure 3B), the peak at 215 nm was probably caused by the conformation change of the P43 peptide into cross-β amyloids. Electron microscopy also showed that fibril structures were formed in the mixture between P43 and NTPs (Supplementary Figure 4E, F). Thus, the propensity of P43 to form cross-β amyloids was probably increased by the interaction with NTPs.

To clarify if RNA binds to the P43 aggregates, we then performed microscopic analysis of the P43 aggregates in the presence and absence of fluorescently-labeled RNA molecules (Figure 2). The P43 peptide (4 mM) formed amorphous aggregations and they appeared the same regardless of the presence of RNA when viewed under bright field. However, fluorescence observation clearly showed that the RNA molecules bound to the P43 aggregates. The interaction between P43 and RNA is probably not dependent on RNA sequence or structure because the tested RNA molecules had different sequences from the target RNA used in the phage display and had no strong secondary structure. When we compared the fluorescent signals from 1–5 nt RNAs (FAM-R1, FAM-R3, FAM-R5), the signal increased as the size of RNA increased, which indicates the P43 peptide can interact with longer RNA molecules more stably. However, when we compared 5–11 nt RNAs (FAM-R5, FAM-R8, FAM-R11), the fluorescent signal rather decreased as the size of RNA increased. This is probably because the surface of the P43 aggregates was saturated with RNA in the case of 5 nt or longer RNAs and longer RNA molecules have fewer fluorescent labels per their covering surface. Interestingly, when we mixed P43 and RNA at high MgCl_2_ concentrations (200–800 mM), the fluorescent signals were still detectable but became blurry (Figure 2), indicating RNA molecules could still be captured by the peptide aggregates but their mobility was increased in such conditions. Furthermore, the addition of NTPs induced visible precipitations of the peptide even at a lower concentration (0.8 mM P43, Supplementary Figure 6, middle), probably by transforming the P43 aggregates into β-amyloid fibrils (Supplementary Figure 3, 4). The resultant P43-NTP co-precipitants could still bind to RNA (Supplementary Figure 6, bottom).

**Figure 2.**
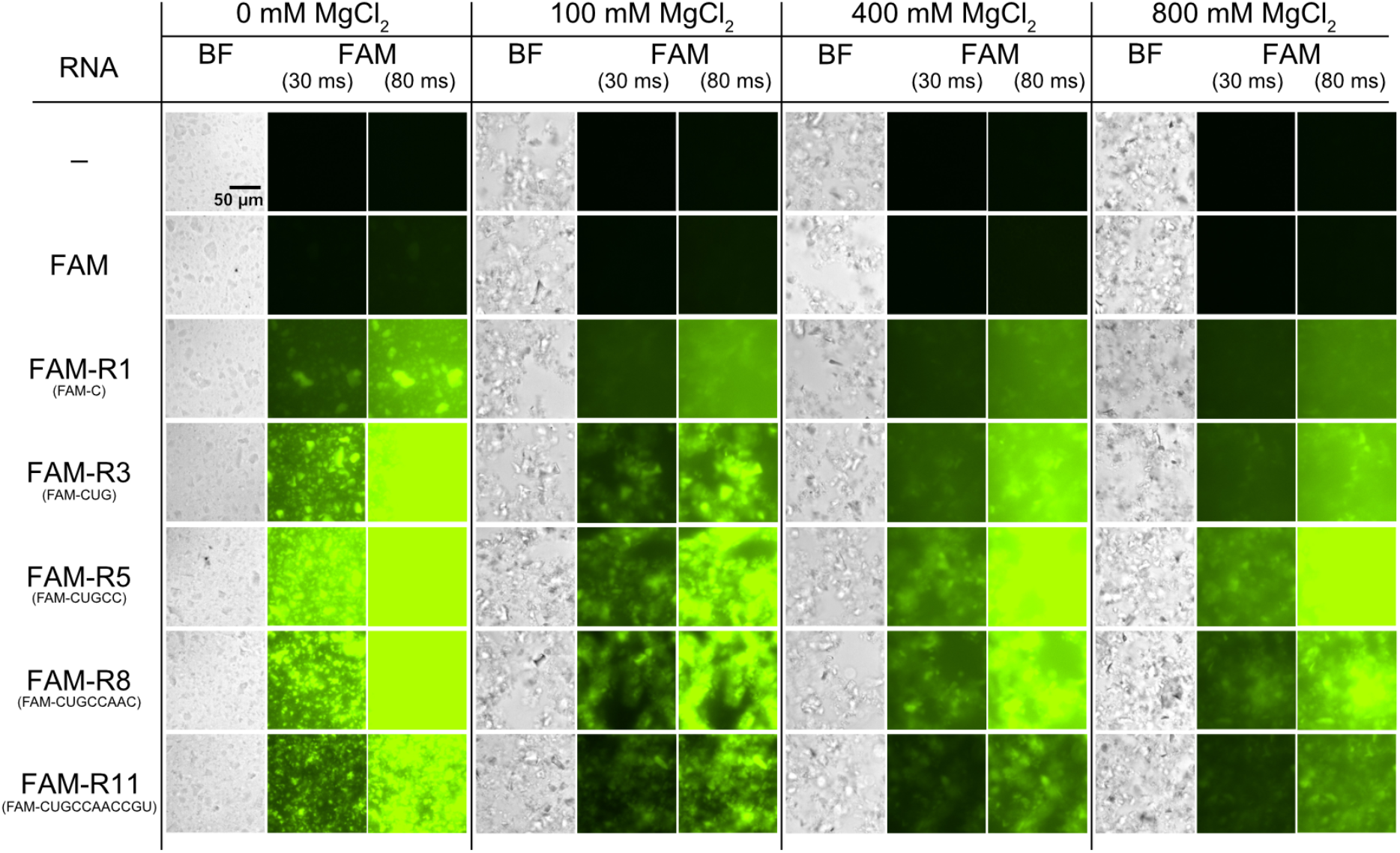
Aggregates of the P43 peptides. Microscopic analysis of the P43 aggregates. The peptide aggregates (4 mM, 2% DMSO) were mixed with fluorescently-labeled RNAs (20 µM FAM-R1–R11) in 0–800 mM MgCl_2_ and observed with bright-field and fluorescence-filter setups (FAM, exposure time = 30 or 80 ms).

### The inhibitory effect of P43 depends on both cationic and hydrophobic amino acid residues

To identify the essential parts of the P43 peptide for its inhibitory effect, we tested truncated variants of P43 at 25 mM MgCl_2_ (Figure 3A). Although the N-terminal and C-terminal amino acid residues fixed in the phage display library (Ala1, Gly9, Gly10, and Ser11) were dispensable for the inhibitory effect (Figure 3A, P43N1, P43C2, and P43C3), further truncation completely abolished the efficacy (Figure 3A, P43N2, P43N4, and P43C4), indicating both of the cationic part (Lys2–Lys3) and hydrophobic part (Val4–Met8) of P43 are essential.

**Figure 3.**
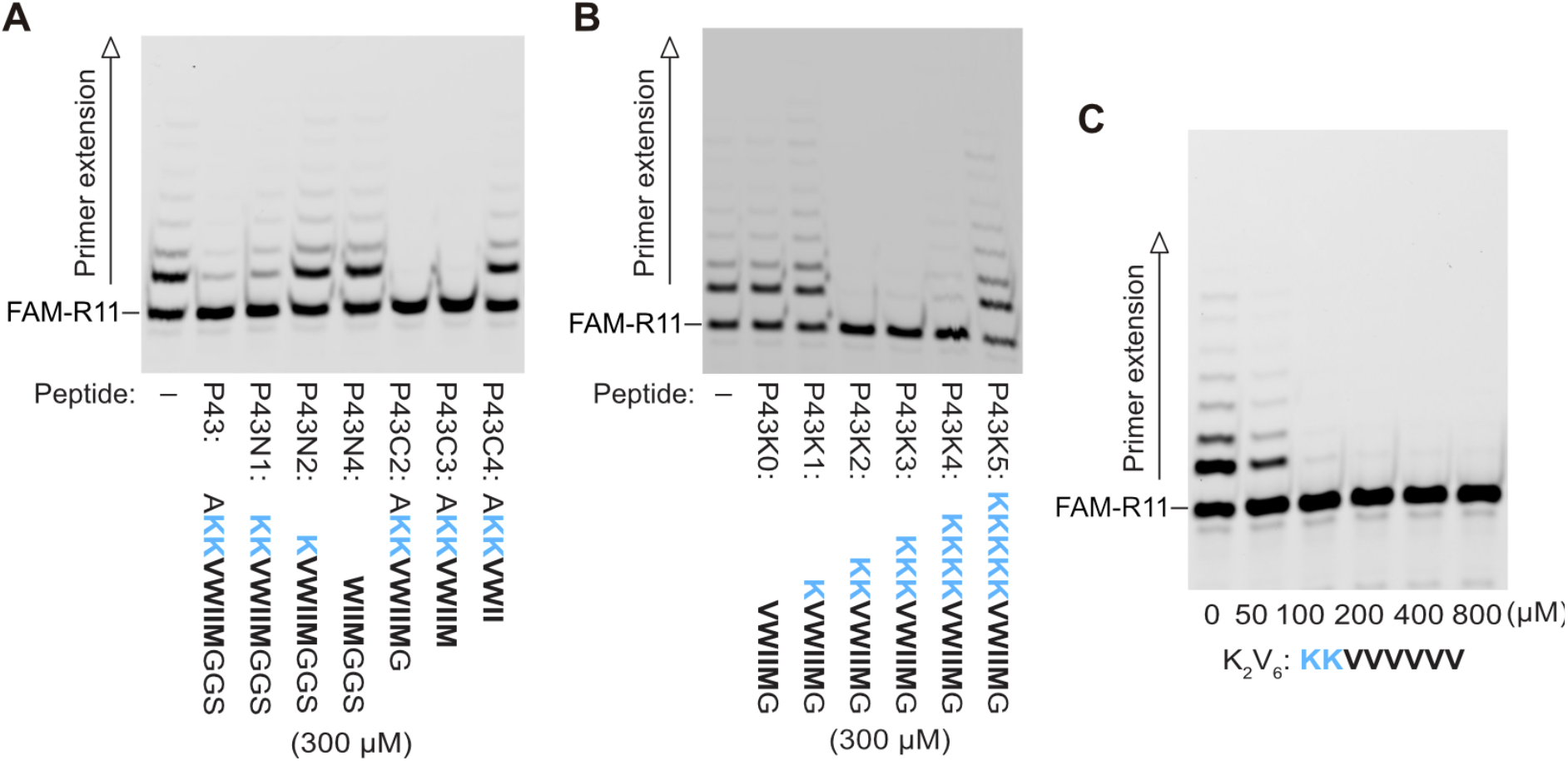
Inhibition of the RPR activity by the P43 variants. Primer extension by RPR Z^LT^ was performed in 50 mM Tris•HCl (pH 8.3) buffer containing 25 mM MgCl_2_, 8% PEG6000, 0.4% (A, B) or 0.8% (C) DMSO and 0.5 mM of each NTP. The reactions were incubated at 17°C for 7 days. P43N4 and P43K0 were completely insoluble and used as 300 µM suspension.

As the interaction between RNA and P43 seems to be mostly electrostatic and achieved by the cationic part (Lys2–Lys3), we then tested P43 variants with different numbers of lysine residues at 25 mM MgCl_2_ (Figure 3B). Consistent with the truncated variants (Figure 3A), the P43 variants with zero or only one lysine residue showed no inhibitive effect on RPR (Figure 3B, P43K0, and P43K1). Although we expected the P43 variants with more than two lysine residues (P43K3, P43K4, P43K5) would interact and inhibit RPR more strongly than the variant with the same number of lysine residues to the original peptide (P43K2), the inhibitory effect rather decreased as the number of lysine residues increased from two (Figure 3B and Supplementary Figure 7). 300 µM P43K5 did not show any inhibitive effect, while 300 µM P43K2–P43K4 inhibited the activity of RPR strongly (Figure 3B). When we reduced the concentration of the peptides to 100 µM, the strong inhibitory effect was observed only with P43K2 (Supplementary Figure 7). Thus, not only the electrostatic interactions between the peptide and RNA molecules but also the hydrophobicity of the peptides are important factors for the inhibitory effect. This observation also indicates P43 works in the aggregated form.

Moreover, we tested another peptide with a simpler hydrophobic-cationic sequence (K_2_V_6_: KKVVVVVV) to investigate if the inhibitory effect on the RPR activity is a general feature of such peptides (Figure 3C). In the presence of 100 µM or more K_2_V_6_, the RPR activity was completely inhibited at 25 mM MgCl_2_, indicating such an inhibitory efficacy like P43 would reside in a wide variety of peptides comprised of hydrophobic and cationic moieties. K_2_V_6_ seemingly also functioned in an aggregated form like P43 as it precipitated when mixed with NTPs, although K_2_V_6_ was apparently soluble in an aqueous stock solution (Supplementary Figure 8).

### The P43 peptide facilitates RNA synthesis by RPR in high salt conditions

As the interaction between RNA and P43 seemed to be weaker in high salt conditions (Figure 2), we speculated if its inhibitory effect on the RPR activity might also be milder in such conditions and performed the RNA primer extension reaction by RPR in buffers containing 200 and 400 mM MgCl_2_ (Figure 4A, B). Surprisingly, in these conditions, the P43 peptide rather enhanced the activity of RPR. At 200 mM MgCl_2_, 400 µM P43 increased the RPR activity roughly three times, while a higher amount of the peptide (800 µM) almost canceled the effect. At 400 mM MgCl_2_, the RPR activity was increased more largely (∼8-fold) in the presence of 400–800 µM P43. We could also observe stimulation of the RPR activity by P43 at 200–800 mM KCl (Supplementary Figure 9). Thus, inhibition of the RPR activity by P43 was probably caused by too tight interactions between RNA and the peptide, which could be canceled in the high salt conditions. Instead, the P43 aggregates stimulated the RPR activity in high Mg^2+^ conditions, probably by concentrating RPR on their surfaces without distorting the active conformation of RPR. It is also interesting that the hydrophobic-cationic P43 peptide could still work in high salt conditions where the purely cationic oligolysine peptide lost its efficacy.^13^ Probably, adopting amyloid conformation increased the multivalency of the cationic charges and maintained weak interactions with RNA even in such high salt conditions. The peptide with a simper sequence (K_2_V_6_) also enhanced RNA synthesis by RPR at 400 mM MgCl_2_ (Figure 4C), indicating such stimulative efficacy would also reside in a wide variety of hydrophobic-cationic peptides.

**Figure 4.**
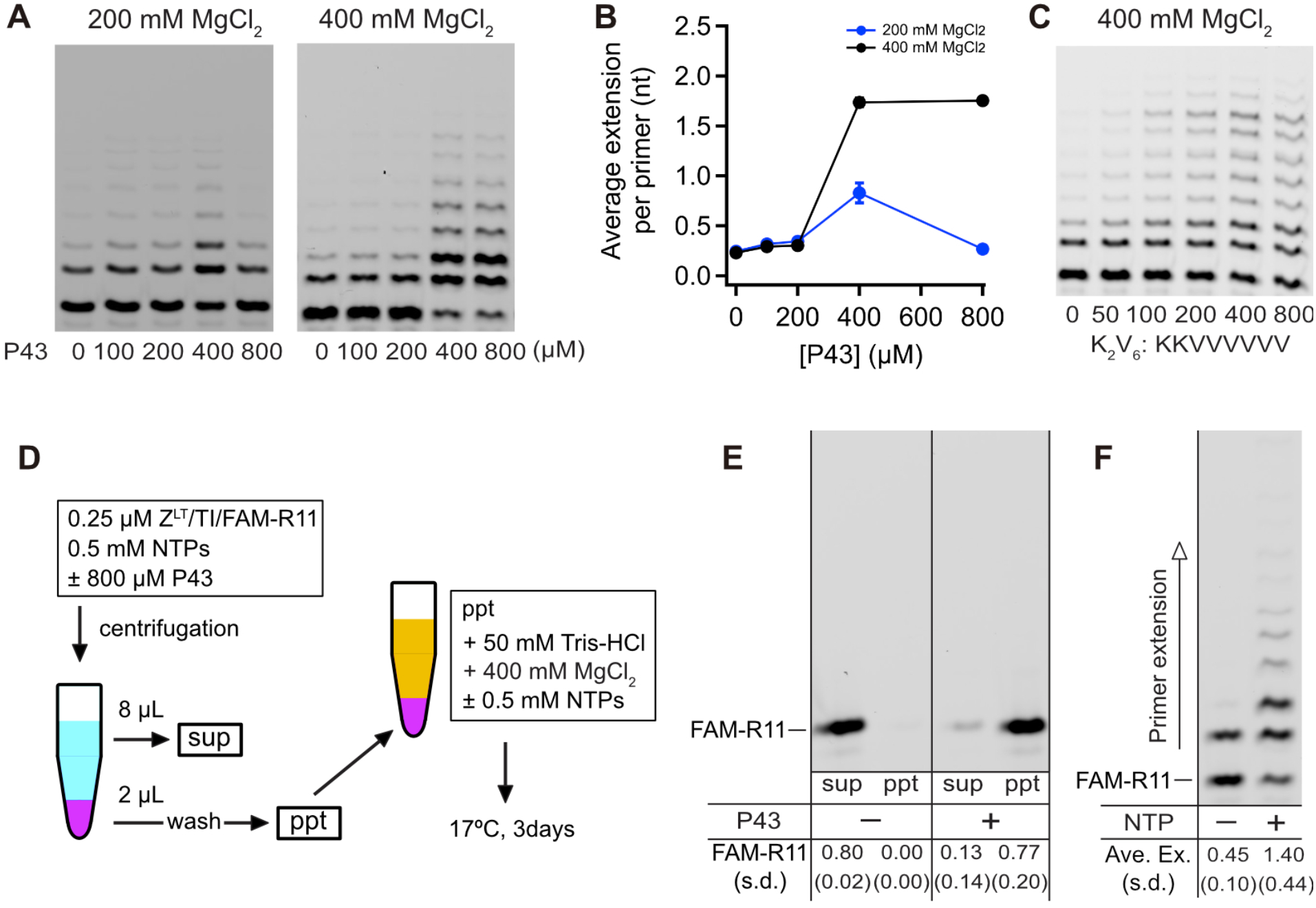
Stimulation of the RPR activity by the P43 peptide in high MgCl*_2_* conditions. (A–C) Primer extension by RPR Z^LT^ was performed in 50 mM Tris•HCl (pH 8.3) buffer containing 200–400 mM MgCl_2_, (A, B) 0.4% or (C) 0.8 % DMSO and 0.5 mM of each NTP. The reactions were incubated at 17°C for 3 days. (B) Error bars represent S. D. (N = 3). (C–E) Co-precipitation and reactivation of RPR with the P43 aggregates. (D) Schematic depiction of the experimental procedure. (E) The fluorescently-labeled primer contained in the supernatant and precipitation samples. The amount of FAM-R11 in the supernatant without P43 was normalized as 0.80. (F) Co-precipitation of RPR and P43 was incubated in 50 mM Tris•HCl (pH 8.3) buffer containing 400 mM MgCl_2_, ± 0.5 mM of each NTP at 17°C for 3 days.

Then, we tested if RPR captured on the P43 aggregates in a low salt condition could be reactivated by resuspension in a high salt buffer (Figure 4D). First, we prepared RNA-peptide premixtures containing RPR, the RNA template/primer, and NTPs in the presence and absence of 800 µM P43. Next, we centrifuged the mixtures to divide them into supernatants and precipitations. Then, we gently washed the precipitations with H_2_O three times. The fluorescently labeled RNA primer was precipitated with the peptide in the presence of P43 (Figure 4E, right), while it was left in the supernatant in the absence of P43 (Figure 4E, left). Finally, we resuspended the P43-RNA precipitations in a reaction buffer containing 400 mM MgCl_2_ with or without additional 0.5 mM NTPs and incubated them for 3 days at 17°C. In the case of the reaction buffer with additional NTPs, the average primer extension by RPR was 1.40 ± 0.44 nt (Figure 4F right), which was comparable to the RPR activity in the same condition without the centrifugation/resuspension processes (the average primer extension at 400 mM MgCl_2_ and 800 µM P43 was 1.75 ± 0.02 nt, Figure 4A, B). These results indicate most RPR was captured on the P43 aggregates and reactivated in the high Mg^2+^ buffer. Even when we resuspended the co-precipitants in the reaction buffer without additional NTPs, RPR showed a detectable activity (Figure 4F, left), although it was significantly weaker than the case with additional NTPs. Absorbance at 260 nm and 280 nm indicated only ∼10% (0.05 mM) of NTPs was co-precipitated with the peptide (Supplementary Figure 3B) in contrast to most RNA molecules were captured on the P43 aggregates (Figure 4E). The difference in absorbance efficiency between NTPs and longer RNAs on such hydrophobic-cationic peptide aggregates might have worked as a selection system for longer RNA molecules on the prebiotic earth. Thus, hydrophobic-cationic peptide aggregates can accumulate RNA in a size-dependent manner on their surface in low salt conditions and reactivate/enhance ribozyme activities in high salt conditions. Such fluctuation between low/high salt conditions would have occurred repeatedly in the dry-wet cycles on the prebiotic earth and also have supported the prebiotic synthesis of biopolymers.^34–40^ Alternatively, the diurnal freeze-thaw cycles would have provided fluctuation of salt concentrations. In eutectic water ice phases, the concentration of Mg^2+^ can be increased without causing quick degradation of RNA, leading to the higher activity of RPR.^41,42^ Hydrophobic-cationic aggregates would have supported the evolution of RNA-based systems in the freeze-thaw cycles, as P43 also enhanced the RPR activity in eutectic ice phases (Supplementary Figure 10).

### Efficacy of P43 is dependent on the sizes of ribozymes

As the P43 peptide was shown to interact with RNA molecules non-specifically (Figure 2), we also investigated its effects on the activities of a few different ribozymes (Figure 5 and Supplementary Figure 11–14). First, we tested the activity of the ribozyme component of RNase P (377 nt), which cleaves the 5′-extension of pre-tRNA to produce matured tRNA.^43^ In this experiment, we prepared an RNA helix mimicking the 5′-extension and the acceptor stem of pre-tRNA (pATSerUG) as the substrate for RNase P (Figure 5A).^44,45^ In low MgCl_2_ condition (25 mM), the activity of RNase P to cleave the 5′-extension of pATSerUG was inhibited in the presence of 400–800 µM P43 in a similar manner to RPR (Figure 5B, Supplementary Figure 11A). Then, we investigated the RNase P activity in higher MgCl_2_ conditions (Figure 5C, Supplementary Figure 11B–D). P43 strongly inhibited the RNase P activity even at 200 mM MgCl_2_. At 400 mM MgCl_2_, P43 still inhibited RNase P ribozyme but in a more moderate way, leaving a significant activity in the presence of 400–800 µM P43. At 800 mM MgCl_2_, P43 finally lost the significant inhibitory effect, indicating the interaction between RNase P and P43 was weakened in the condition. The efficacy of P43 on RNase P was not significantly influenced in the presence of NTPs (Supplementary Figure 12). These results were in stark contrast to the case of RPR, which was rather activated by P43 in the high salt conditions (Figure 4). This is probably because RNase P (377nt) is about twice longer than RPR Z^LT^ (221 nt) and unproductive interactions between RNase P and P43 can occur even in 200–400 mM MgCl_2_.

**Figure 5.**
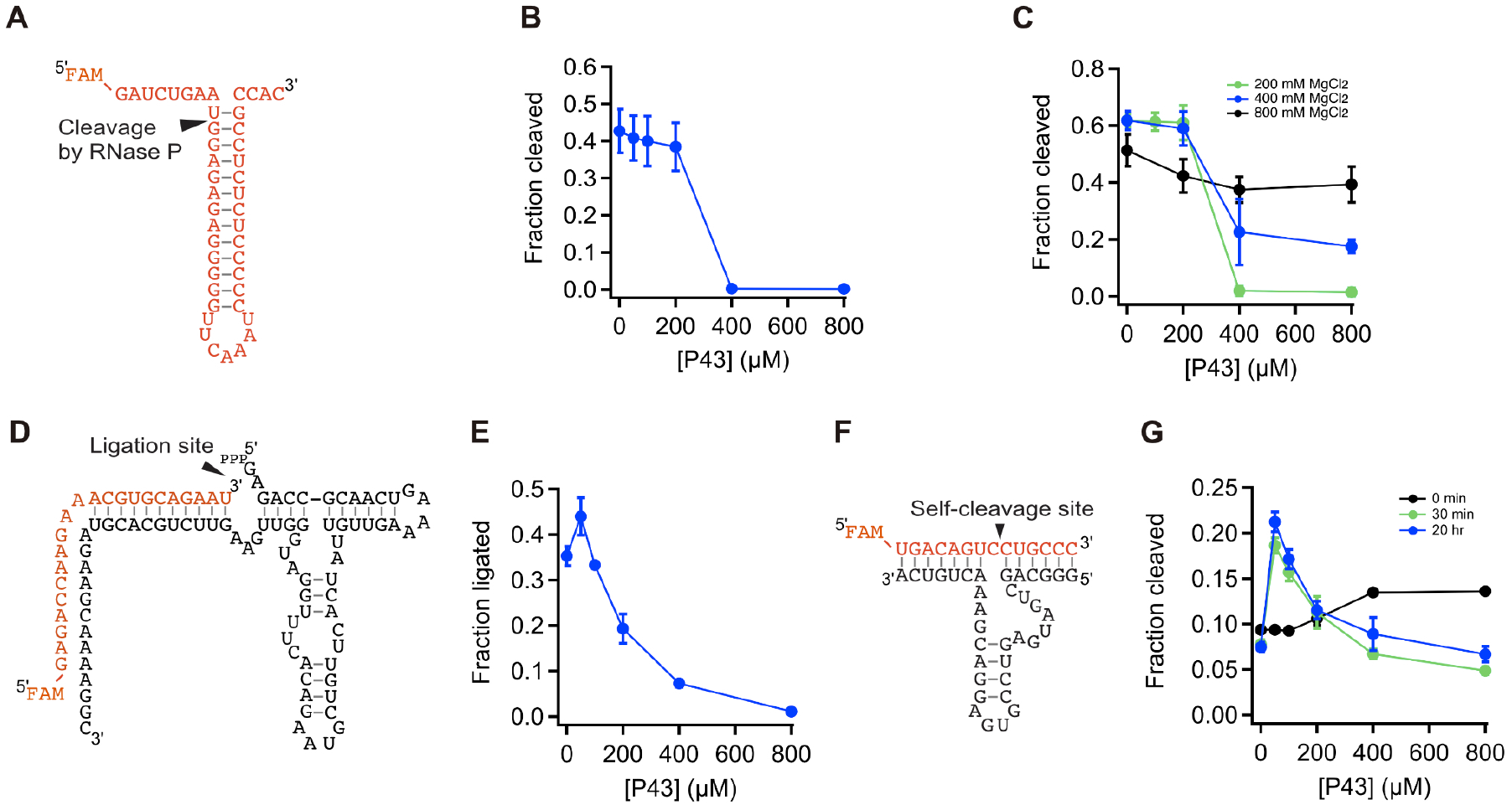
Activity of various ribozymes in the presence of P43. (A) Schematic depiction of the substrate for the RNase P ribozyme (pATSerUG). (B–C) The substrate cleavage reaction by RNase P in the presence of P43 at different MgCl_2_ concentrations. The reactions were incubated (B) in 50 mM Tris•HCl (pH 8.0) buffer containing 25 mM MgCl_2_ at 37°C for 10 min, or (C) in 50 mM Tris•HCl (pH 8.0) buffer containing 200–800 mM MgCl_2_ at 37°C for 2.5 min. (D) Schematic depiction of F1* ligase. (E) The self-ligation reaction by F1* ligase was performed in 50 mM Tris•HCl (pH 8.3), 25 mM MgCl_2_, 0.4%DMSO at 25 °C for 20 sec. (F) Schematic depiction of trans-cleaving hammerhead ribozyme. (G) The substrate cleavage reaction by hammerhead ribozyme in 50 mM Tris•HCl (pH 8.3), 25 mM MgCl_2_, 0.4%DMSO. The ribozyme and peptide were preincubated for 0 min (black), 30 min (green), or 20 hours (blue) before the reactions were started by addition of MgCl_2_. The reactions were incubated at 25 °C for 20 sec. (B, C, E, G) Error bars represent S. D. (N = 3).

Then, we also tested the efficacy of P43 on the activity of a shorter ribozyme, F1* ligase ribozyme (Figure 5D, 81 nt ribozyme strand + 22 nt substrate strand),^46^ assuming the interactions between the shorter ribozyme and P43 are weaker than the interactions between longer ribozymes (RPR and RNase P) and P43. We found that even in a low MgCl_2_ condition (25 mM), the self-ligation activity of F1* ligase was not completely inhibited by 400 µM P43 (Figure 5E, Supplementary Figure 13), while the activities of RPR and RNase P were abolished in the same condition (Figure 1C, 5B). These observations indicate that most ribozymes can be inhibited when captured on such peptide aggregates in the low salt condition. Longer ribozymes (e.g., RPR, RNase P) tend to be more severely affected by P43 than shorter ribozymes (e.g., F1* ligase) probably because longer ribozymes are structurally more flexible and can be easily distorted from their active conformation when they are stuck on the surface of peptide aggregations.

Finally, we tested the activity of an even shorter ribozyme, trans-cleaving hammerhead ribozyme (Figure 5F, 35 nt ribozyme strand + 14 nt substrate strand)^47^ in the presence of the P43 peptide. When we first premixed the peptide with the reaction buffer containing Mg^2+^ and then mixed them with ribozymes to start the reaction, the ribozyme activity was slightly enhanced in the presence of 400–800 µM P43. (Figure 5G, 0 min preincubation, Supplementary Figure 14, left). Speculating the reaction time of the hammerhead ribozyme (20 sec) was too short for the peptide to interact with such a short ribozyme, we then premixed hammerhead ribozyme and the P43 peptide and incubated for 30 min or 20 hours before the reactions were started by adding the reaction buffer. In these conditions, the P43 peptides more largely stimulated the activity of the hammerhead ribozyme (Figure 5G, 30 min and 20 hr preincubation, Supplementary Figure 15, middle and right). The ribozyme activity was increased about 2.5-fold when 100 µM P43 was mixed, while the stimulative effect was gradually canceled when the concentration of the P43 was further increased. As the structure of the hammerhead ribozyme is most simple among the tested ribozymes, its active structure may be stable enough even when it is stuck on the peptide aggregates in the low Mg^2+^ condition. The enhancement of the activity of hammerhead ribozyme by P43 is probably caused by the concentration of the RNA molecules on the peptide surface, as similar concentration and enhancement effect for hammerhead ribozyme in LLPSs was reported previously.^27,48^

## CONCLUSION

Here, we have reported that a new RNA binding peptide, P43 (AKKVWIIMGGS) selected by phage display, forms amorphous aggregates and influences the activity of various ribozymes. In the low salt condition (25 mM MgCl_2_), the P43 peptide strongly inhibited RNA primer extension by RPR (Figure 1C, D). Both cationic (Lys2-Lys3) and hydrophobic (Val4–Met8) parts of the peptide was required for the inhibitory effect (Figure 3A, B), indicating the cationic amino acid residues are essential for the interaction with RNA, while the hydrophobic residues are required to cause aggregation of the peptide. 5 nt or longer RNA molecules can bind to the surface of the P43 peptide aggregates stably (Figure 2). At high concentrations of MgCl_2_ (≥ 200 mM), the interactions between RNA and the peptide were still detectable but became weaker. In such high salt conditions, the P43 peptide rather enhanced the activity of RPR (Figure 4A, B), probably concentrating RNA molecules on their surface without strong distortion of the ribozyme structure. Furthermore, RPR captured on the peptide aggregates in a low salt condition (0 mM MgCl_2_) could be reactivated by resuspension in a high salt condition (400 mM MgCl_2_) (Figure 4D–F). The observations that a simpler peptide (K_2_V_6_) showed similar regulatory effects support the possibility that such hydrophobic-cationic peptides have existed on the prebiotic earth. (Figure 2C, 4C)

These results also suggest the potential roles of such hydrophobic-cationic peptide aggregates in the emergence of the RNA-based life on the prebiotic earth (Figure 6). Wet-dry cycles have been suggested to be critical for the prebiotic synthesis of biopolymers including peptide and RNA.^34–40^ as polymerization reactions without complicated chemistry for substrate activation are basically dehydration reactions between monomers. In such wet-dry cycles, the environment surrounding the synthesized polymers should have inevitably fluctuated and the biopolymers needed to survive such a variety of conditions to be finally integrated as life. Peptides and their analogs could be easily synthesized by the dry-down condition^34–38^ and the hydrophobic peptides would remain as aggregates without dissipating when the inflow of water brought various materials from different environments. In the wet/diluted conditions, cationic peptide aggregates like P43 could have accumulated scarce nutrition and RNA molecules. This process would also have served as the selection for longer RNA molecules as longer RNA apparently bound to the P43 peptide more stably (Figure 2, 4E). When the aqueous solution was partially evaporated and condensed, such peptide aggregates could have enhanced ribozymes like RPR as the concentrations of salt and nutrition were increased. Such condensed conditions could have also supported biopolymer formation.^38–40^ In this way, the wet-dry cycles with cationic peptide aggregates could have enriched longer and functional RNA molecules. Alternatively, such fluctuation of salt and nutrient concentrations would have occurred in diurnal freeze-thaw cycles, in which eutectic ice phases could also have stimulated the function and evolution of RPR.^41,42^ Once the concentration of RNA molecules had reached high enough, more elaborate systems for compartmentalization like LLPSs would have emerged. Therefore, peptide aggregates could have supported the emergence of life in harsh prebiotic environments before the first cell-like structure had established.

**Figure 6.**
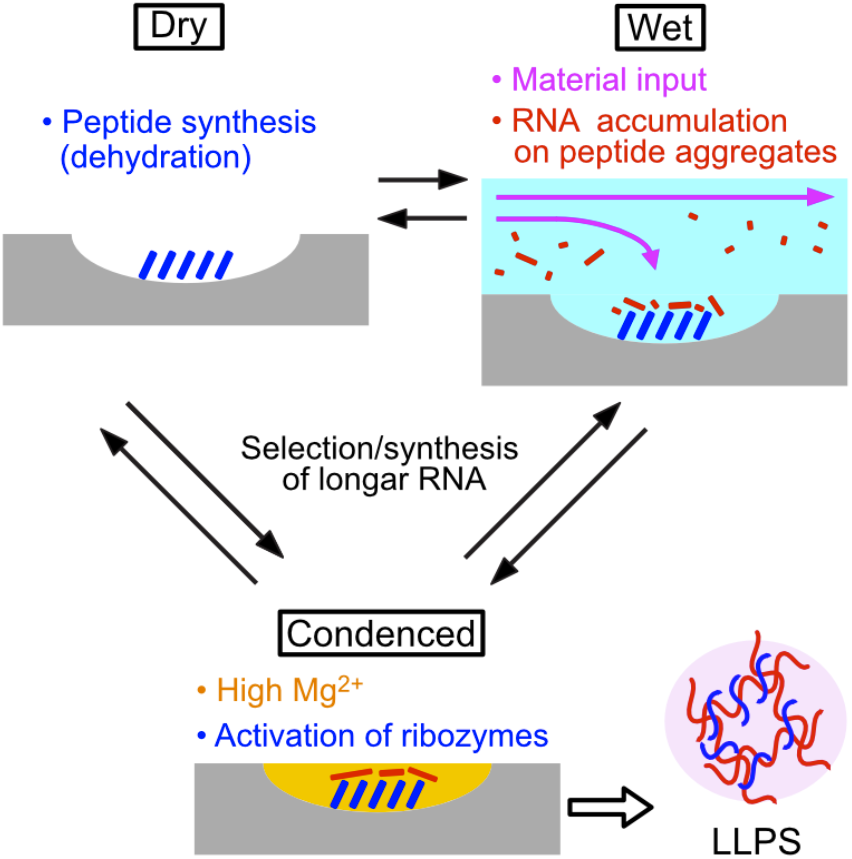
Potential roles of peptide aggregates in the wet/dry (freeze/thaw) cycles. Once peptide aggregates (blue) formed in the dry phases in wet-dry cycles, they would have stably existed and captured nucleic acids (red) in the wet phases. Then, after evaporation (freezing) of water, the peptide aggregates would have enhanced activities of ribozymes in condensed conditions with high [Mg^2+^].

## Supporting Information

*Supporting Information Available*: This material is available free of charge *via* the Internet.

Methods and Supplementary Figures 1–14 (PDF)

## Acknowledgements

We thank Stefan Luzi for assistance in the phage display experiment. We also thank Tomomi Uchikubo-Kamo for helping the electron microscopic analysis. The authors are grateful to the beamline staff scientists at KEK. P.L. and S.T were supported by JSPS (20K14599 and 18H01328). S.T. was also supported by the Astrobiology Center of National Institutes of Natural Sciences (AB301005).

## Supporting Information

### METHODS

#### Preparation of RNAs and peptides

The DNA and RNA sequences used in this study is listed on Supplementary Table 1 and 2. Z^LT^, Zc and RNase P ribozymes are synthesized by in vitro transcription as previously described.^1–3^ Then, the 3′-end of Zc was biotinylated by using the periodate oxidation method.^4^ Briefly, the Zc was incubated in 66 mM sodium acetate (pH 4.5), 5 mM NaIO_4_ on ice for 45 min in the dark. After the incubation, the ribozyme was purified by iso-propanol precipitation. Then, Zc was incubated in 20 mM sodium acetate (pH 6.1), 0.2% SDS, 70% DMSO, 7 mM biotin-LC-hydrazid (Thermo Fisher Scientific) overnight at 23°C in the dark. Finally, the labeled Zc ribozyme was purified by ethanol precipitation.

The sequence for F1* ligase was amplified by PCR using gF1*_C, gF1*_F, and gF1*_R as template and primers. Then 5′-triphosphorirated L1* ligase was transcribed in an in-vitro transcription kit (MEGAshortscript T7 transcription kit, Invitrogen). Chemical synthesis and HPLC purification of hammerhead ribozyme and fluorescently-labeled RNA substrates (FAM-R1–11, pATSerUG, substrate strands for F1* ligase and hammerhead ribozyme) were performed by Japan Bio Services. pATSerUG was further gel-purified to remove contaminants.

All peptides tested in this report were supplied by Japan Bio Services. We dissolved the peptides in 100% DMSO and then diluted with H_2_O to prepare stock solutions.

#### Peptide selection

To select peptides that bind to RPR, we applied the previously-reported phage display method,^5^ omitting the chemical cyclization procedure of the peptide libraries. We performed 3–4 rounds of biopanning of the peptide library with seven randomized amino acid residues (AXXXXXXXGGS) against the biotinylated Zc immobilized on magnetic beads. Then, we sequenced ∼30 and ∼50 phagemids after the third and fourth rounds of biopanning, respectively. The sequences without any stop codon are listed in Supplementary Figure 1B.

#### Activity test of RNA polymerase ribozyme

Initially, RPR Z^LT^ was annealed with the fluorescently labeled RNA primer (FAM11) and RNA template (TI) in H_2_O (50 °C for 5 min, 17 °C for 10 min). Then, the ribozyme complex was mixed with the extension buffer, so that each reaction contained 0.25 µM ZLT/FAM11/TI, 50 mM Tris•HCl (pH 7.6 or 8.3), 25–400 mM MgCl_2_, 0 or 8% PEG6000, 0.4%DMSO, 0.5 mM of each NTP, and the peptides. The samples were incubated at 17 °C for 3 or 7 days. PEG6000 was mixed in the low [Mg^2+^] reactions as molecular crowding agent to increase the reaction rate.

Then, the primer extension reactions were stopped by adding 3–6 volumes of the stop buffer (8 M urea, 80 mM EDTA, and 10 µM TC2). After heat treatment (94°C, 5 min), the samples were resolved by urea-PAGE gels (20% poly-acrylamide, 8M urea). The gels were analyzed using an Amersham Typhoon scanner (GE Healthcare).

#### Activity test of RNase P ribozyme

First, *E. coli* RNase P ribozyme and pATSerUG were annealed in 1 mM EDTA (pH 8.0), by heating the samples at 70 °C for 5 min, and gradually decreasing the temperature to 12 °C. Then, the ribozyme complex was mixed with the reaction, so that each reaction contained 0.25 µM RNase P/pATSerUG, 50 mM Tris•HCl (pH 8.0), 25–800 mM MgCl_2_, 0.4%DMSO, and the peptides. The samples were incubated at 37 °C for 2.5–10 min. Then, the RNA cleavage reactions by RNase P were stopped by adding 3–12 volumes of the stop buffer (8 M urea, 80 mM EDTA). After heat treatment (94°C, 5 min), the samples were resolved by urea-PAGE gels (10–15% poly-acrylamide, 8M urea). The gels were analyzed using an Amersham Typhoon scanner.

#### Activity test of F1* ligase

F1* ligase was annealed with labeled substrate strand, F1*sub in H_2_O, by heating the samples at 70 °C for 5 min, and gradually decreasing the temperature to 12 °C. Next, the ribozyme complex was mixed with peptide solution and allowed to stand for 30 min at 25 °C. Then, the reactions were started by adding the buffer including Tris•HCl and MgCl_2_, so that each reaction contains 0.25 µM F1*/F1*sub, 50 mM Tris•HCl (pH 8.3), 25 mM MgCl_2_, 0.4%DMSO, and the peptides. The samples were incubated at 25 °C for 20 sec. Then, the reactions were stopped by adding three volumes of the stop buffer (8 M urea and 50 mM EDTA). After heat treatment (94°C, 5 min), the samples were resolved by urea-PAGE gels (10–15% poly-acrylamide, 8M urea). The gels were analyzed using an Amersham Typhoon scanner.

#### Activity test of the hammerhead ribozyme

HH35 ribozyme was annealed with a labeled substrate, HPshortFAM in 1mM EDTA (pH 8.0), by heating the samples at 70 °C for 5 min, and gradually decreasing the temperature to 12 °C. For 0 min preincubation reactions, the ribozyme complex was mixed with the reaction buffer, so that each reaction contains 0.25 µM HH35/HPshortFAM, 50 mM Tris•HCl (pH 8.3), 25 mM MgCl_2_, 0.4%DMSO, and the peptides. For 30 min and 20 hr preincubation reactions, the ribozyme complex was mixed with peptide solution and allowed to stand for 30 min and 20 hr at 25 °C. Then, the reactions were started by adding the reaction buffer. The samples were incubated at 25 °C for 20 sec. Then, the reactions were stopped by adding three volumes of the stop buffer (8 M urea and 50 mM EDTA). After heat treatment (94°C, 5 min), the samples were resolved by urea-PAGE gels (10–15% poly-acrylamide, 8M urea). The gels were analyzed using an Amersham Typhoon scanner.

#### Microscopic observation of P43 aggregates

Aggregate of P43 was observed by inverted microscope system (Nikon, Eclipse Ti) under bright field observation. FAM or FAM-labeled RNAs were added into P43 suspension with MgCl_2_. These samples contained 4 mM P43 (2% DMSO), 20 µM FAM/RNAs and 0–800 mM MgCl_2_. Then, P43/RNA complex solutions of 8 µL were applied to cell counting slides (Bio-Rad) and visualized by using fluorescence setup with GFP filter set (excitation at 470 nm).

#### Negative-stain electron microscopy

Structures of P43 in the absence / presence of NTPs were observed by negative-stain electron microscopy (FEI, TF20). These samples contained 4 mM P43 (2% DMSO) / 0.8 mM P43 (0.4% DMSO), and 0/0.5 mM NTPs. Samples were applied to the carbon film grids, and then stained with 2% uranyl acetate. EM images were taken at (200 kv) a magnification of 50,000.

### Supplementary Tables

**Supplementary Table 1.**
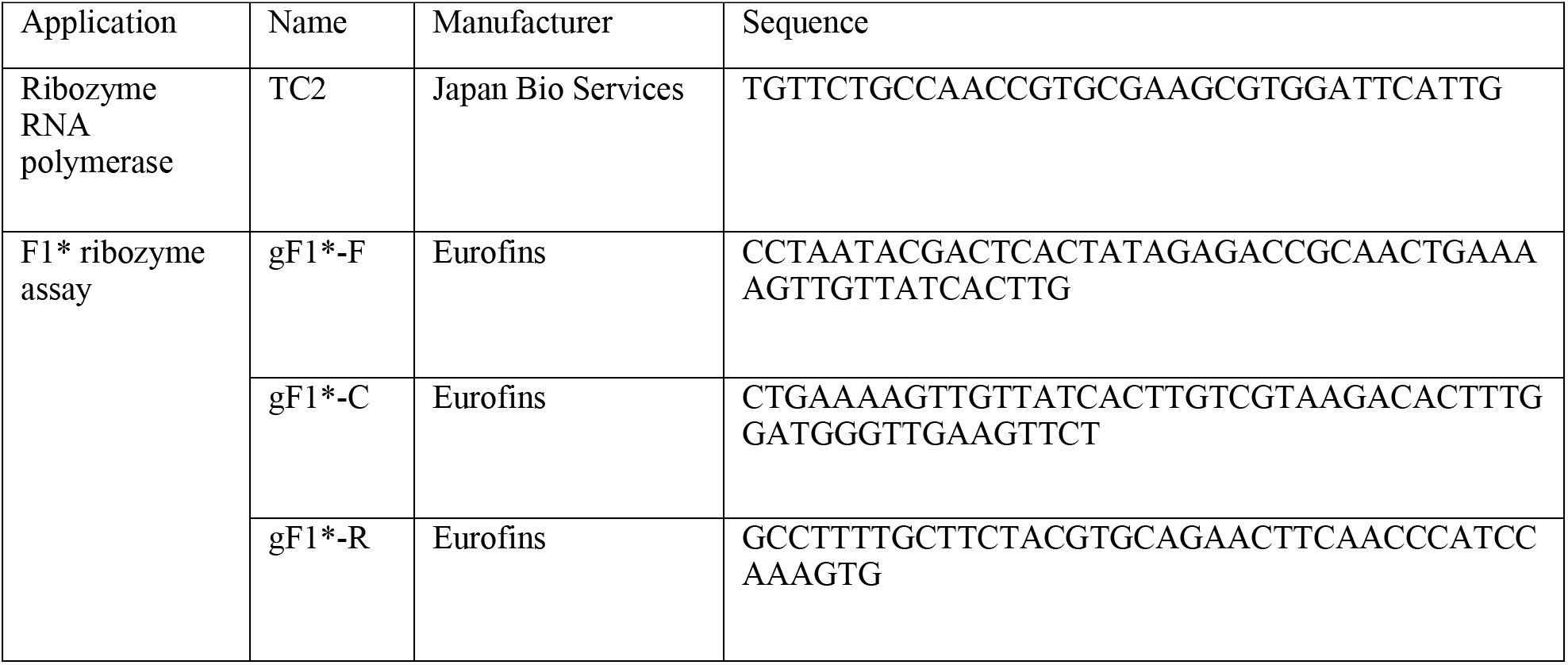
DNA sequences

**Supplementary Table 2.**
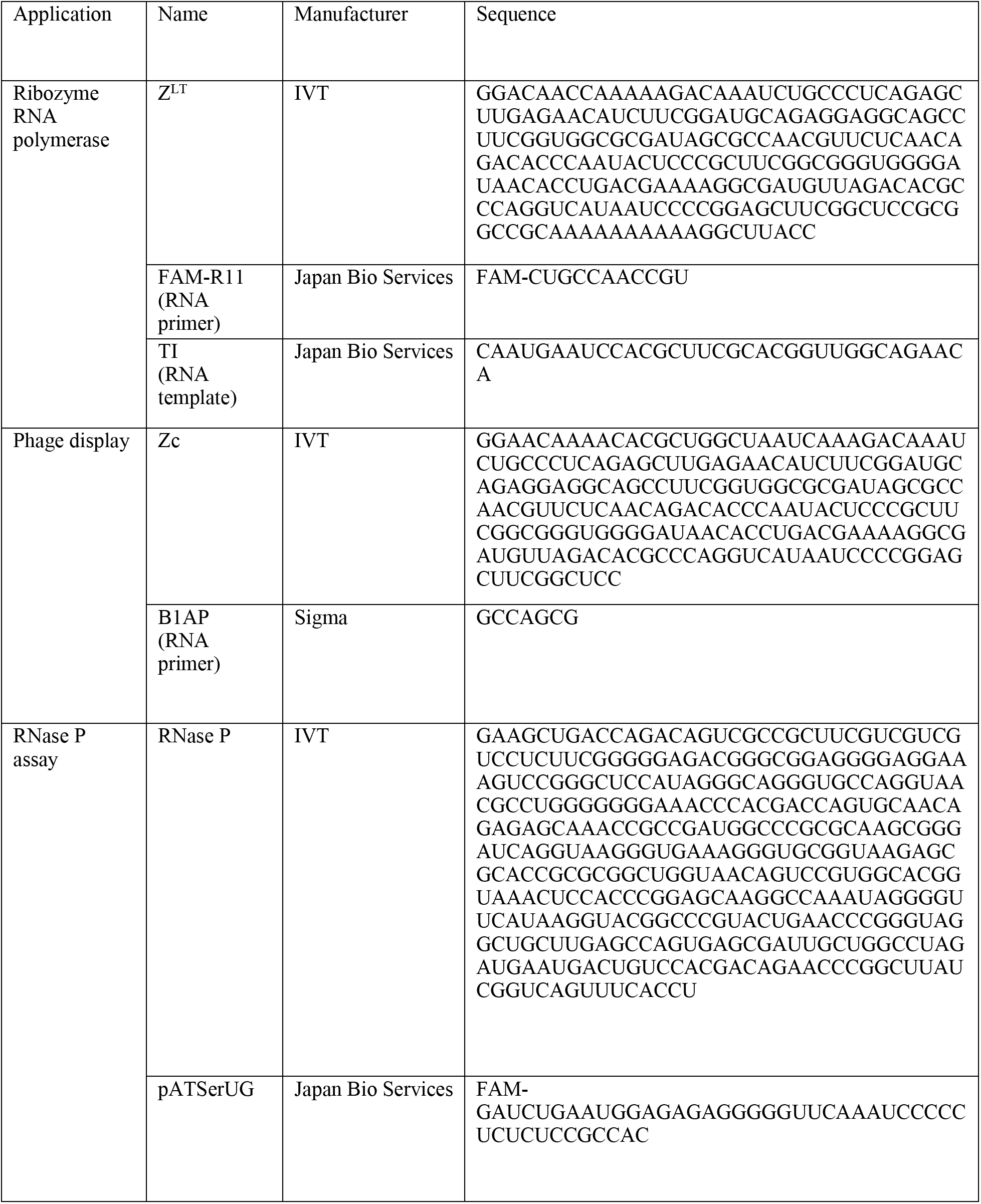
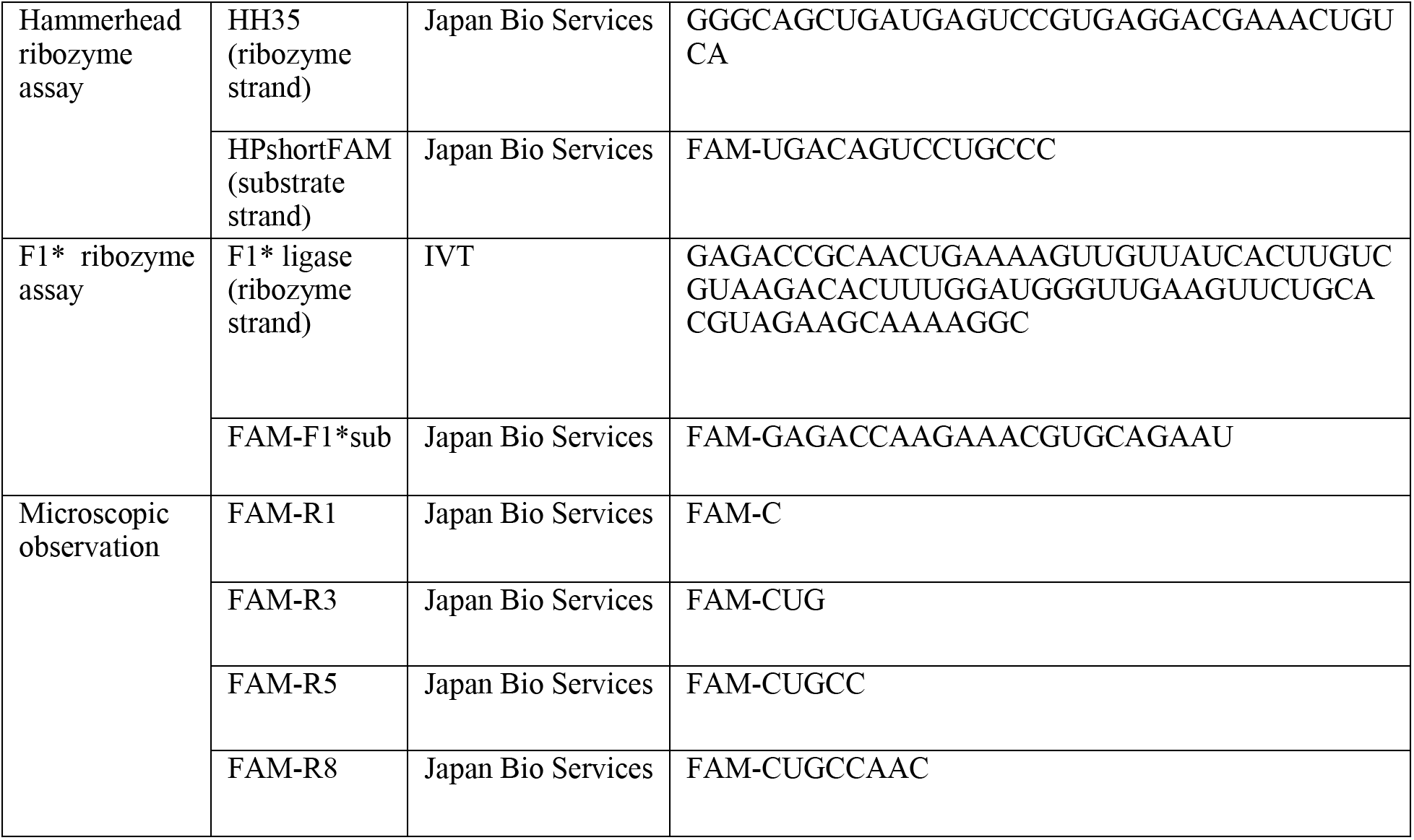
RNA sequences

### Supplementary Figures

**Supplementary Figure 1.**
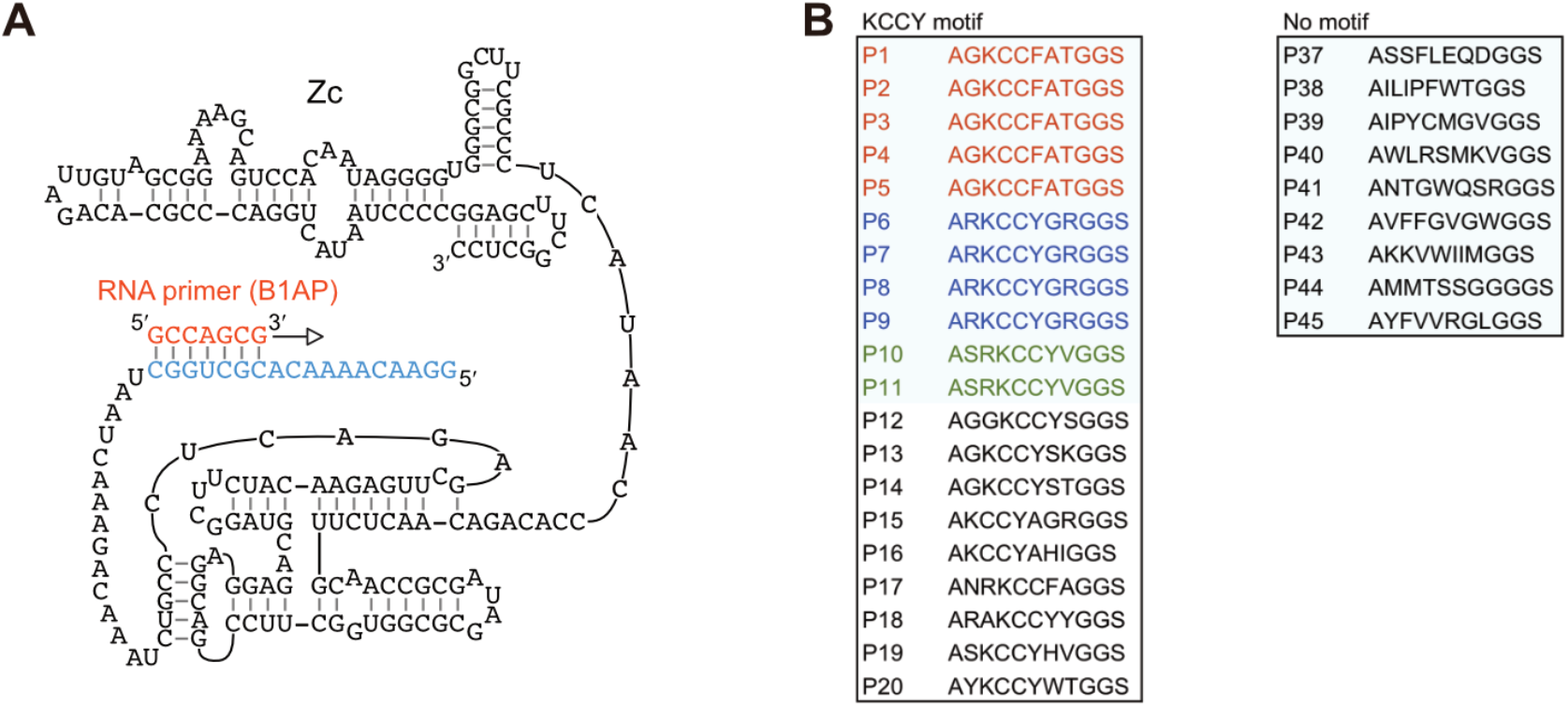
Phage display targeting RPR. (A) The target RNA used in the phage display experiment (Zc). The 3′-biotinylated Zc ribozyme was annealed with the RNA primer (B1AP) in water (50°C for 5 min, 17°C for 10 min) before immobilization on the magnetic beads. (B) Sequences of the selected peptides. Only sequences without any stop codon are shown.

**Supplementary Figure 2.**
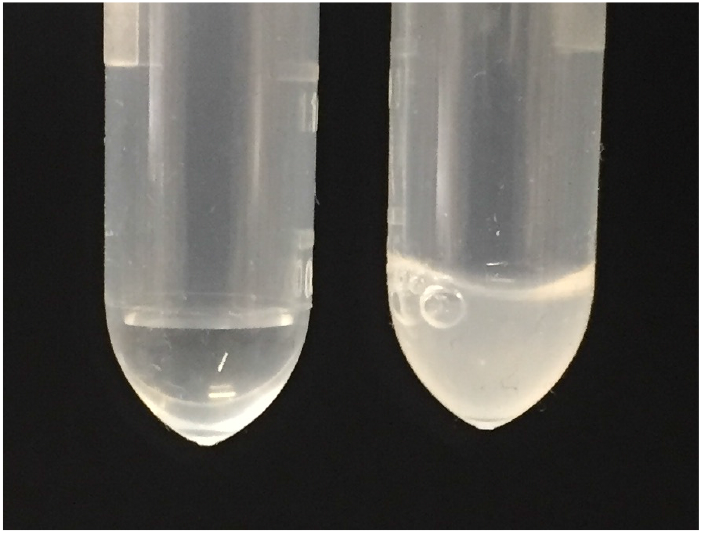
Precipitant of the P43 peptide. Right: Precipitants in the stock solution of P43 (20 mg/mL P43 in 20% DMSO) kept in a freezer for more than one night; Left: H_2_O.

**Supplementary Figure 3.**
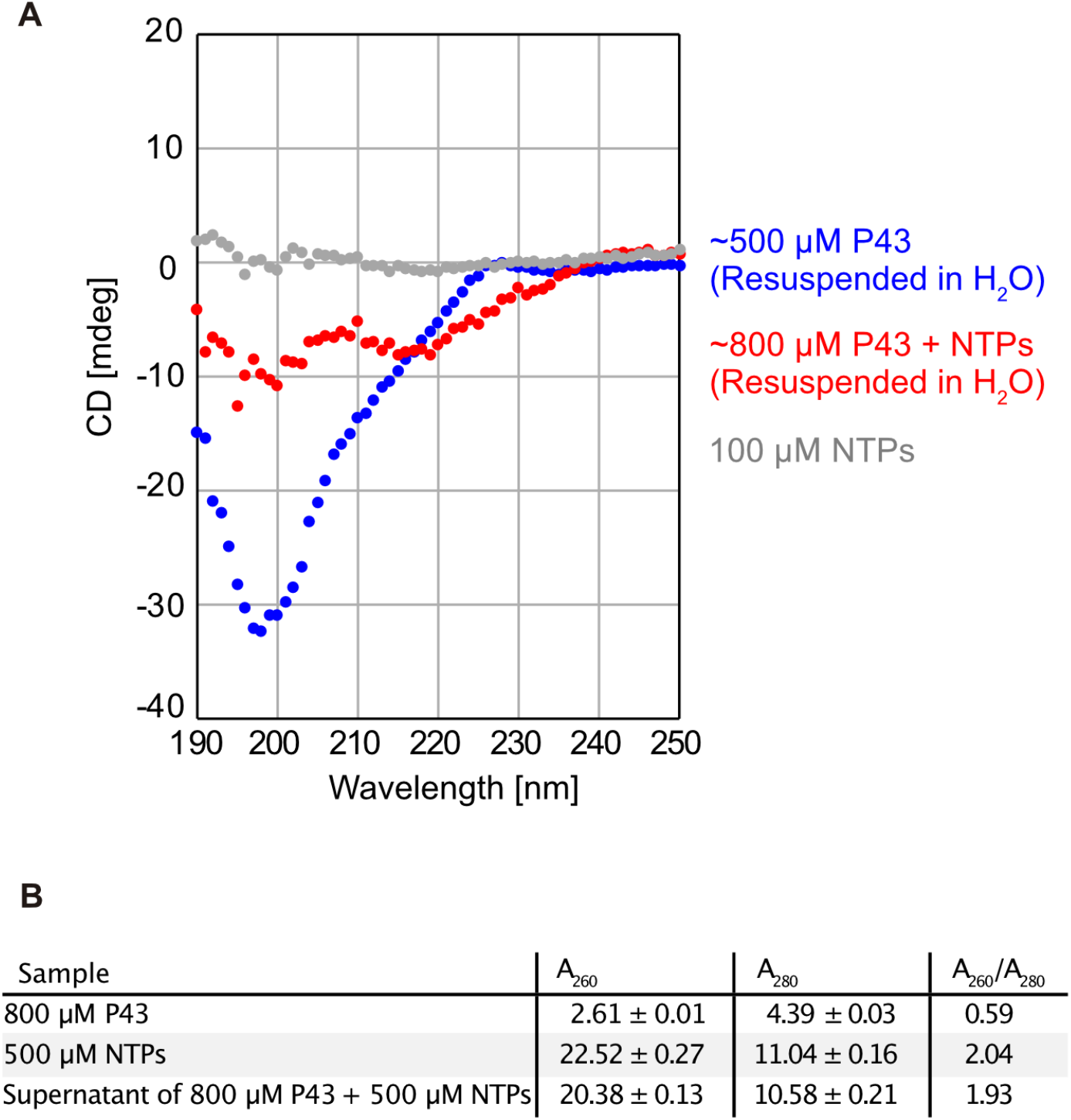
CD spectra of the P43 peptide. (A) The aggregates in the P43 stock solution (20 mg/mL P43 in 20% DMSO) and the solution mixed with NTPs (800 µM P43, 0.5 mM NTPs, and 0.4% DMSO) were precipitated by centrifugation and gently washed with H_2_O 4–5 times to remove DMSO. Then, the P43 aggregates were resuspended in H_2_O. 1 mm-pathlength cuvettes were filled with 200 µL resuspended solution, and the CD spectra were scanned from 250 nm to 190 nm at 20°C using a JASCO J820 CD spectrometer. While the resuspended P43 from the stock solution (500 µM) showed typical CD spectra of random coiled peptides (blue), co-precipitants of P43 and NTPs exhibited another negative peak at 215 nm (red). CD spectra of 0.1 mM NTPs are also shown (gray). (B) Absorbances at 260 nm and 280 nm of the samples were measured using NanoDrop (Thermo Scientific). Roughly 10% of mixed NTPs (0.05 mM) were estimated to be contained in the co-precipitants with P43.

**Supplementary Figure 4.**
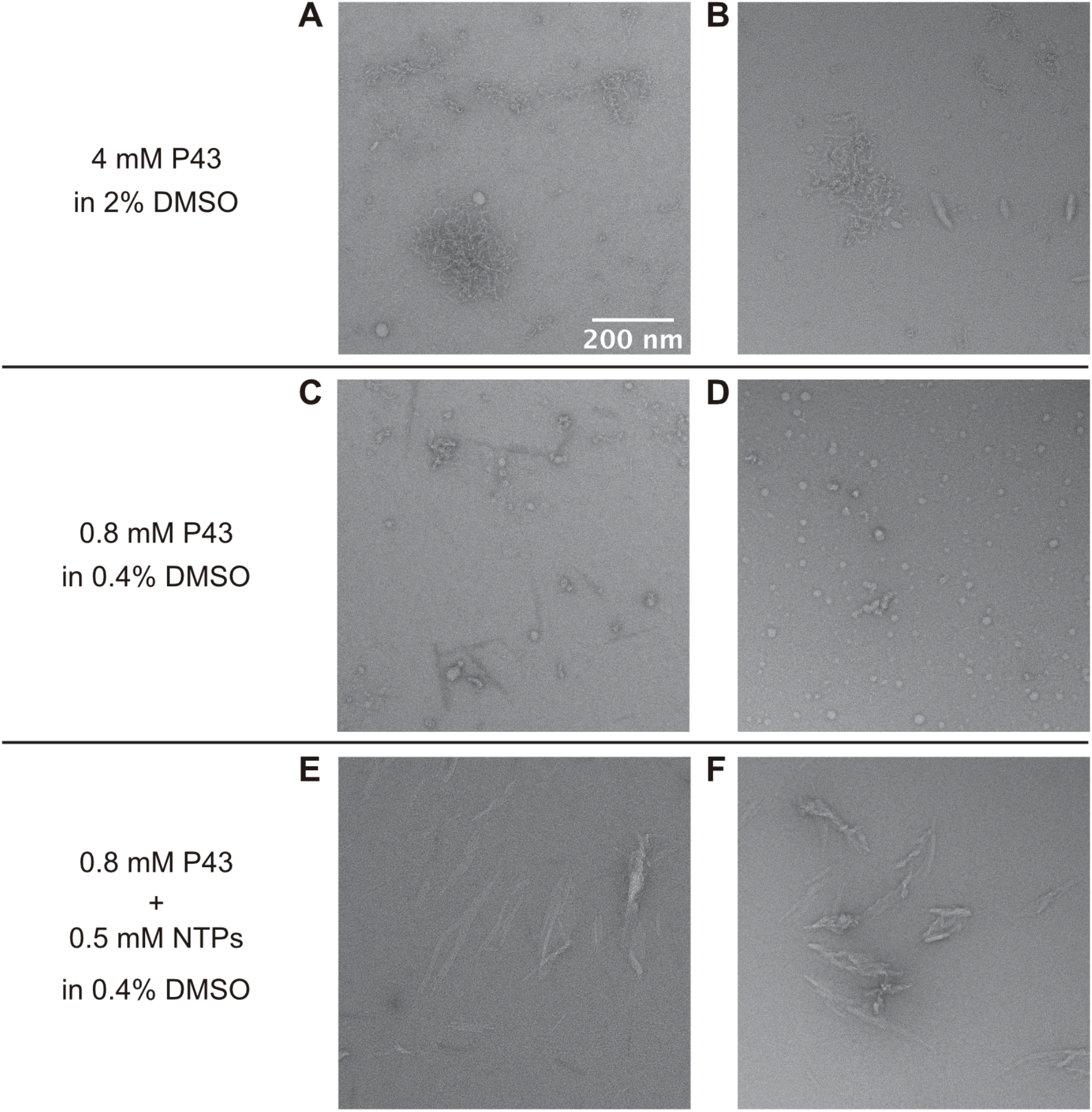
Electron microscopic analysis of the P43 aggregates. (A, B) The P43 aggregates (4 mM P43 in 2% DMSO) were observed by using negative-stain electron microscopy multiple times. (C, D) the P43 aggregates at a concentration (0.8 mM P43 in 0.4% DMSO) used in the activity test of the ribozymes. (E, F) Aggregates of 0.8 mM P43 with 0.5 mM NTPs in 2% DMSO.

**Supplementary Figure 5.**
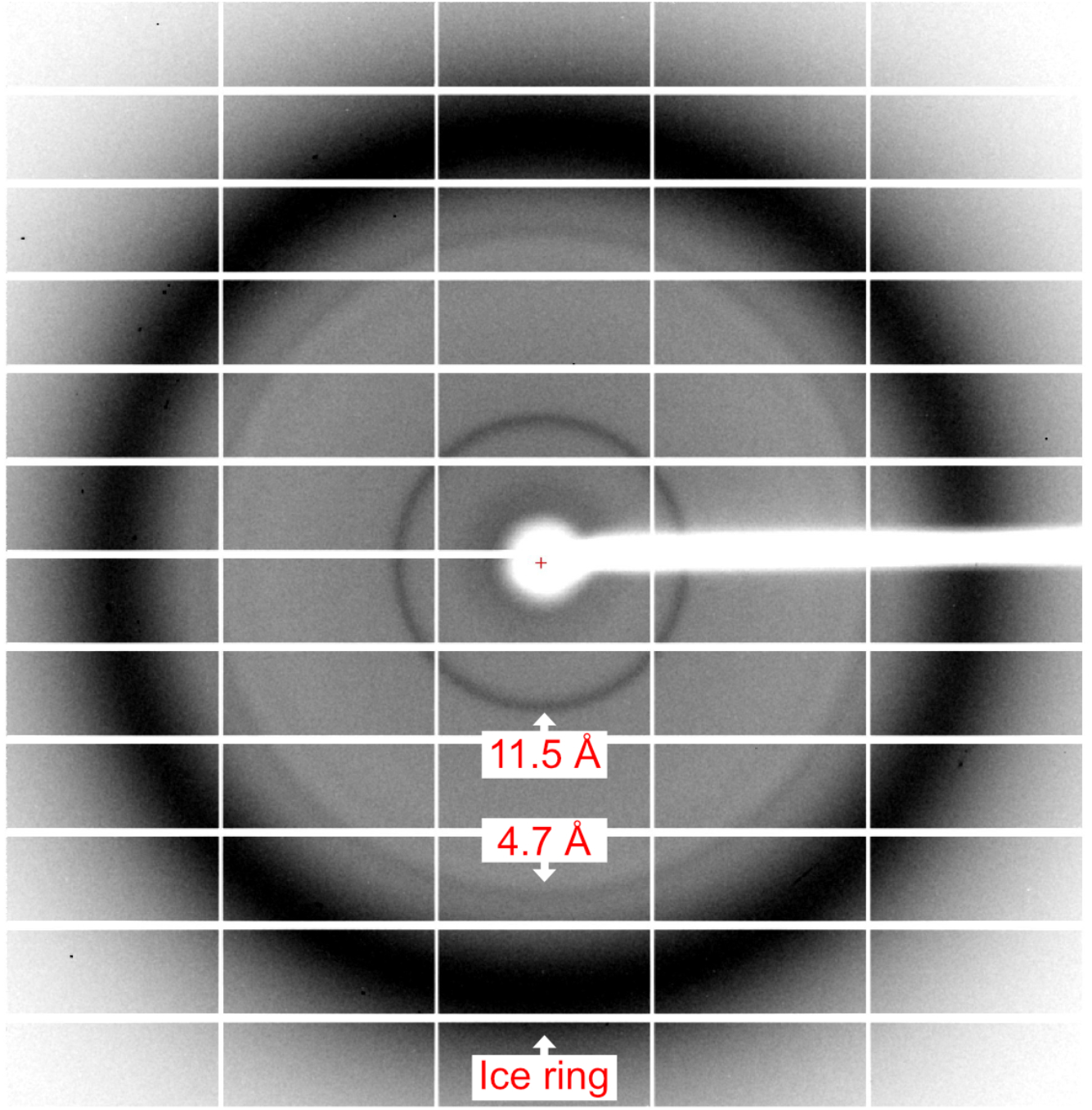
X-ray diffraction data from the P43 aggregates. The peptide aggregates in 10 mg/mL P43 in 10% DMSO was precipitated by centrifugation, picked with a loop for protein crystallography, and frozen with liquid N_2_. Then, the X-ray diffraction data was obtained by synchrotron radiation (KEK, PF BL5A). Diffractions at 4.7 Å and ∼10 Å are general features of cross-β amyloid.^6^

**Supplementary Figure 6.**
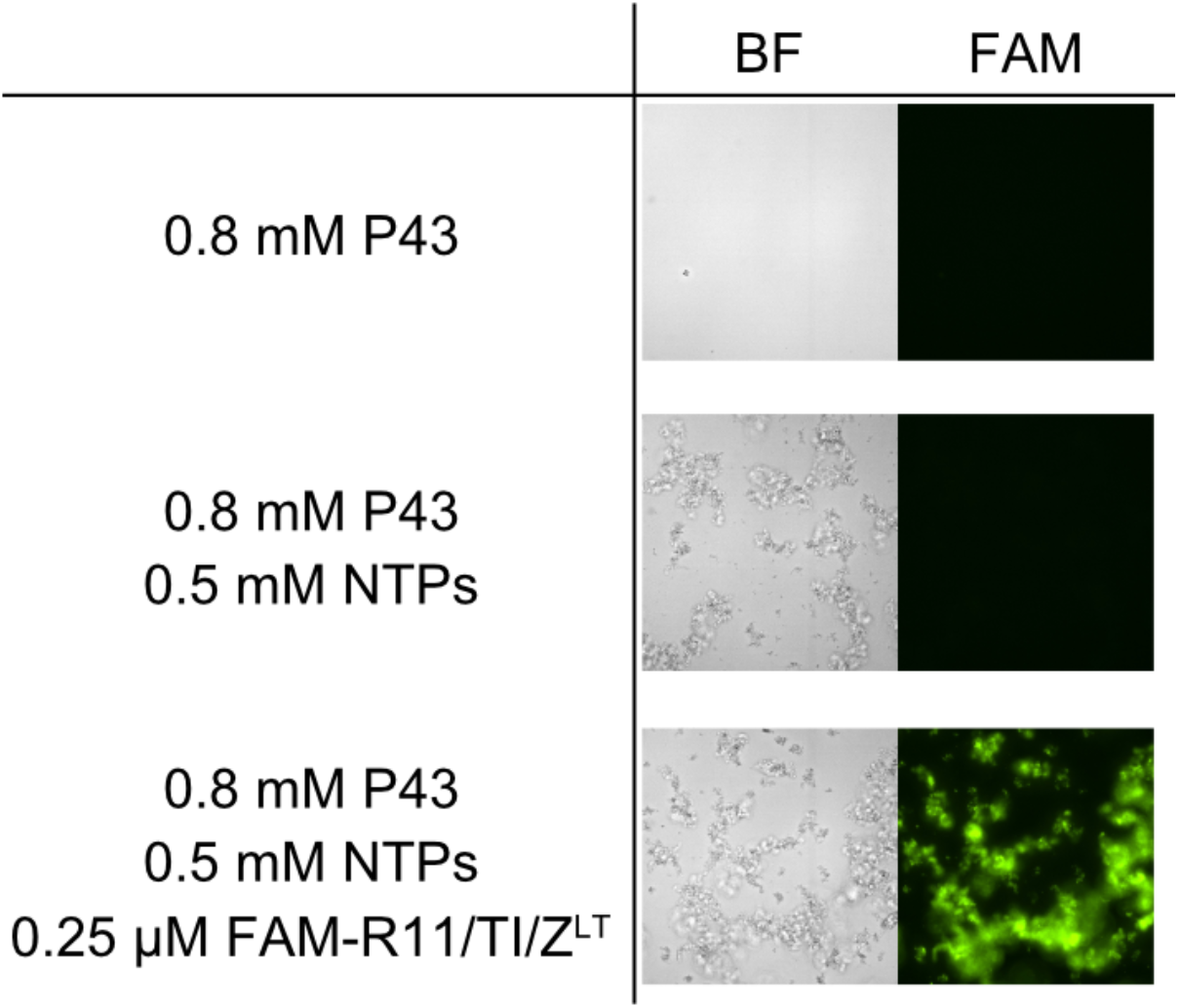
Aggregates of the P43 peptide with NTPs. Microscopic analysis of the aggregates of P43 with NTPs. The peptide aggregates (0.8 mM, 0.4% DMSO) were mixed with the fluorescently-labeled RNA primer, RNA template, and RPR (0.25 µM FAM-R11/TI/Z^LT^) at 0 MgCl_2_ and observed with bright field microscopy and fluorescence filter setup (FAM, exposure time = 80 ms). Without NTPs, the P43 solution was apparently clear, although we could observed some amorphous aggregates in the same condition by electron microscopic analysis (Supplementary Figure 4C, D).

**Supplementary Figure 7.**
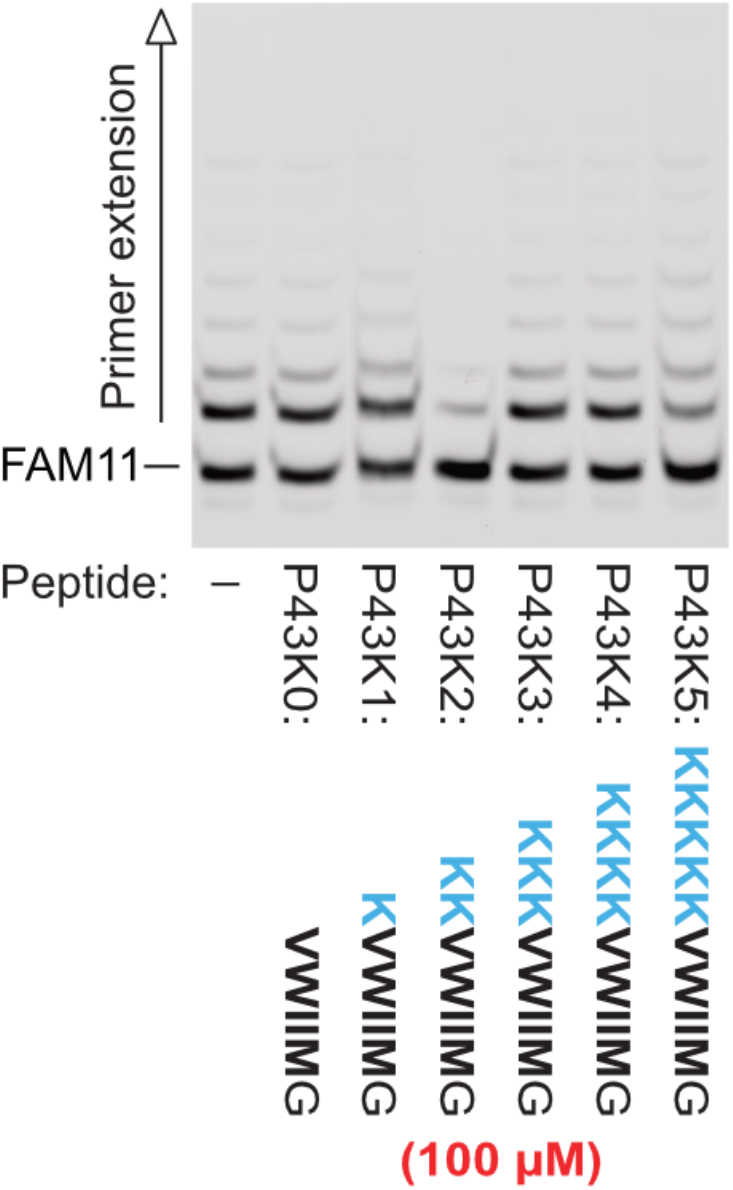
The P43 variants different numbers of lysine residues. Primer extension by RPR Z^LT^ was performed in 50 mM Tris•HCl (pH 8.3) buffer containing 25 mM MgCl_2_, 8% PEG6000, 0.4% DMSO and 0.5 mM of each NTP with 100 µM P43 variants. The reactions were incubated at 17°C for 7 days. P43K0 were completely insoluble and used as suspension. Only P43K2 showed a significant inhibitory effect.

**Supplementary Figure 8.**
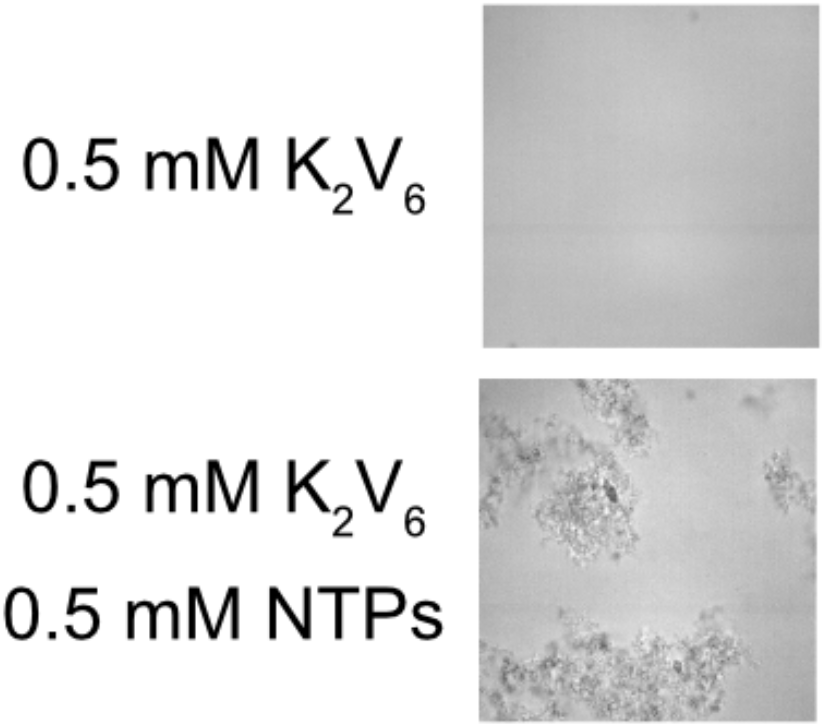
Aggregates of the K*_2_*V*_6_* peptide with NTPs. Microscopic analysis of the aggregates of K_2_V_6_ with NTPs. The peptide solutions with and without NTPs were observed with bright field microscopy.

**Supplementary Figure 9.**
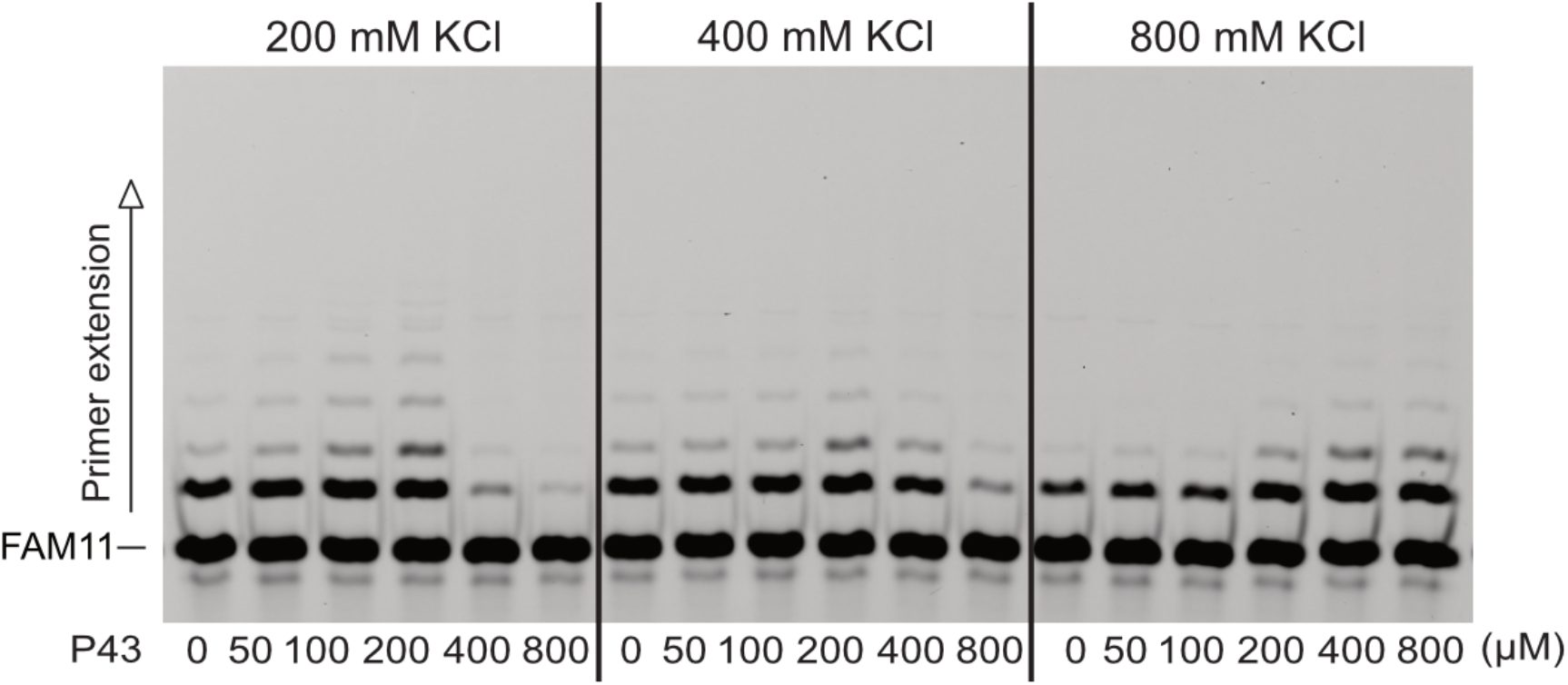
The efficacy of the P43 peptide at high KCl concentrations. Primer extension by RPR Z^LT^ was performed in 50 mM Tris•HCl (pH 8.3) buffer containing 25 mM MgCl_2_, 200–800 mM KCl, 8% PEG6000, 0.4% DMSO and 0.5 mM of each NTP. The reactions were incubated at 17°C for 7 days. Although the RPR activity was slightly inhibited at high KCl concentrations, the stimulative effect of P43 could be still observed.

**Supplementary Figure 10.**
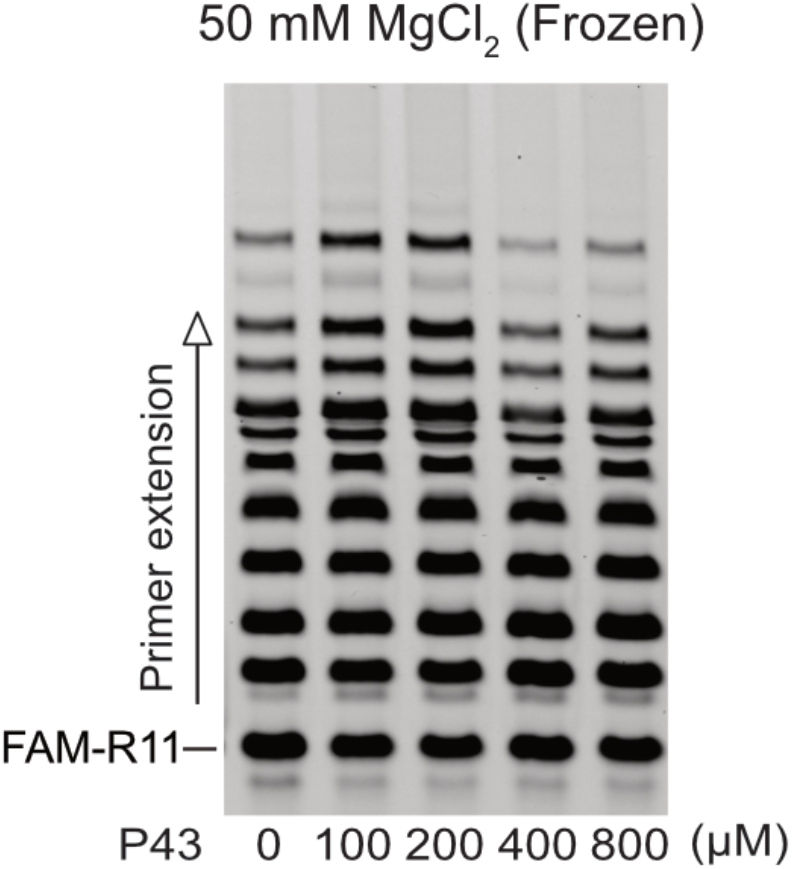
The efficacy of the P43 peptide in eutectic ice phases. Primer extension by RPR Z^LT^ was performed in 50 mM Tris•HCl (pH 8.3) buffer containing 50 mM MgCl_2_, 0.4% DMSO and 0.5 mM of each NTP. The reactions were incubated at −7°C for 7 days after frozen at −80 °C for 30 min. In eutectic ice phases, the increase of the RNA and Mg^2+^ concentrations causes the higher activity of RPR. Further stimulation of the RPR activity could be observed in the presence of 100–200 µM P43.

**Supplementary Figure 11.**
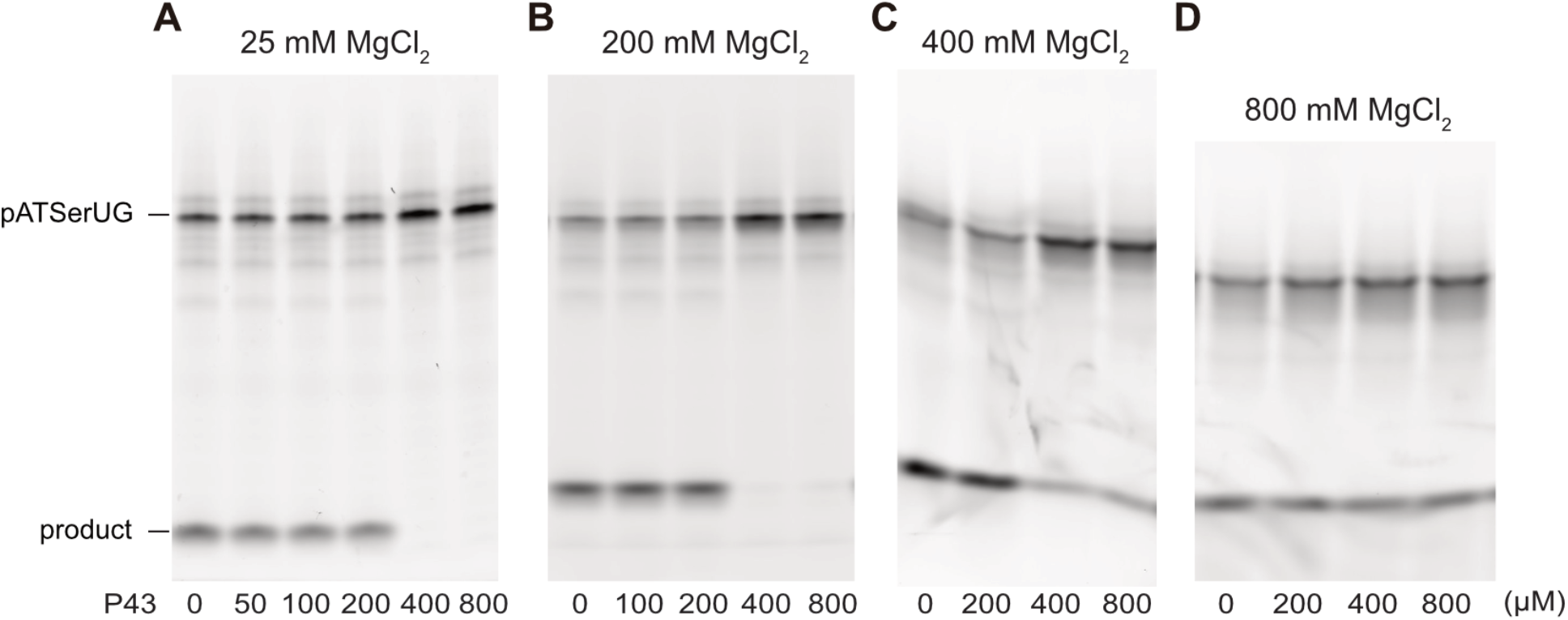
Inhibition of the RNase P activity by the P43 peptide. (A–D) The RNase P activity was tested in the presence of the P43 peptide. The reactions were performed in 50 mM Tris•HCl (pH 8.0) buffer containing 25 mM MgCl_2_ at 37°C for 10 min, (B–D) 50 mM Tris•HCl (pH 8.0) buffer containing 200–800 mM MgCl_2_ at 37°C for 2.5 min.

**Supplementary Figure 12.**
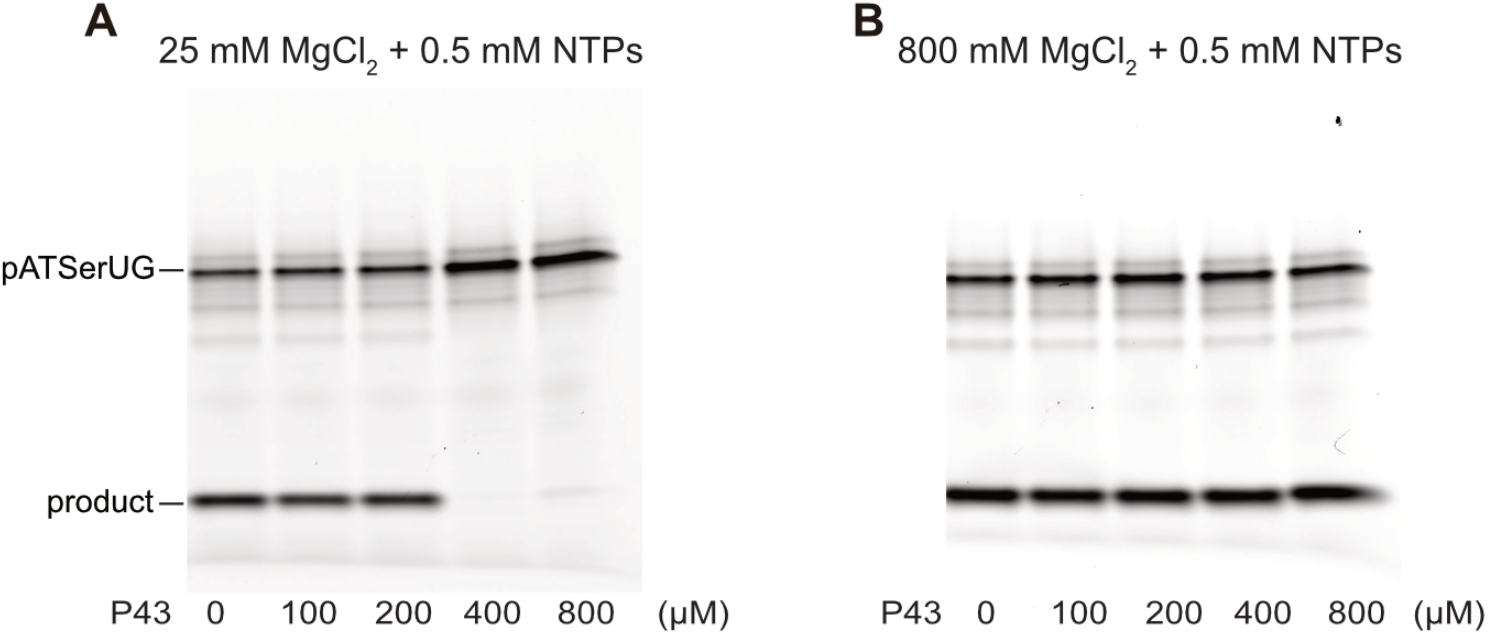
Inhibition of the RNase P activity by the P43 peptide in the presence of NTPs. (A, B) The RNase P activity was tested in the presence of the P43 peptide and NTPs. The reactions were performed in (A) 50 mM Tris•HCl (pH 8.0) buffer containing 25 mM MgCl_2_ and 0.5 mM NTPs at 37°C for 10 min, (B) 50 mM Tris•HCl (pH 8.0) buffer containing 800 mM MgCl_2_ and 0.5 mM NTPs at 37°C for 2.5 min.

**Supplementary Figure 13.**
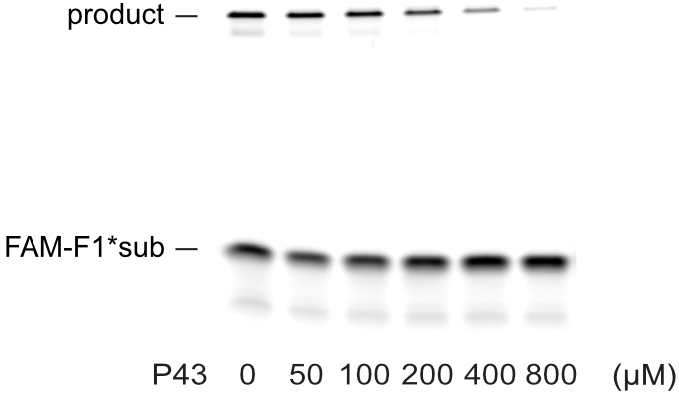
Inhibition of the F1* activity by the P43 peptide. The F1* activity was tested in the presence of the P43 peptide. The reactions were performed in 50 mM Tris•HCl (pH 8.3) buffer containing 25 mM MgCl_2_ at 25°C for 20 sec.

**Supplementary Figure 14.**
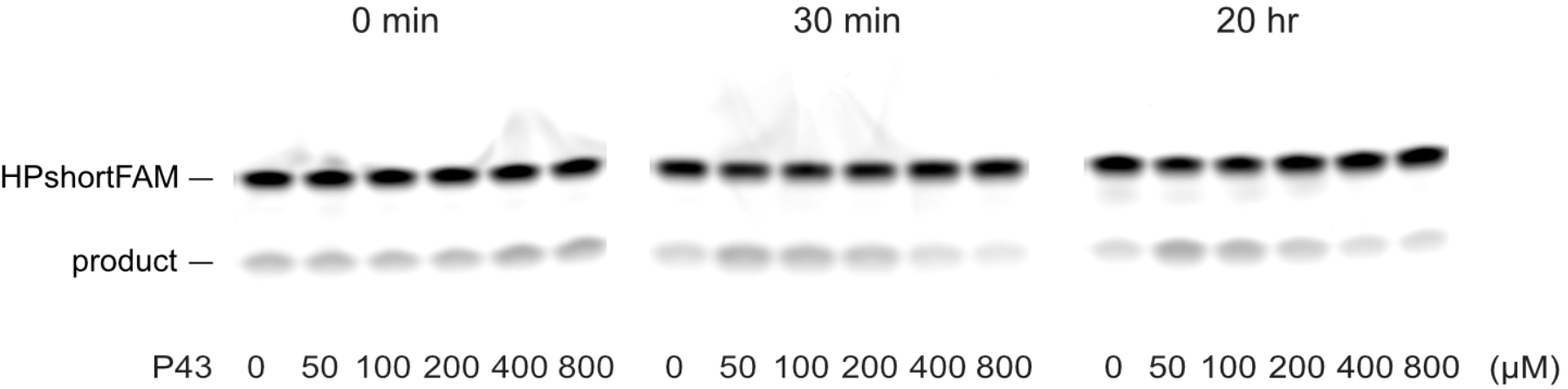
The HH35 activity affected by the mixing time of HH35 and P43 peptide. (A–C) The HH35 activity was tested after 0–20 hr preincubation in the presence of the P43 peptide. The reactions were performed in 50 mM Tris•HCl (pH 8.3) buffer containing 25 mM MgCl_2_ at 25°C for 20 sec.

## Notes

### Competing Interest Statement

The authors have declared no competing interest.

